# Quantifying antibiotic susceptibility and inoculum effects using transient dynamics of *Pseudomonas aeruginosa*

**DOI:** 10.1101/2025.11.26.690803

**Authors:** Sarah Sundius, Jennifer Farrell, Kelly L. Eick, Rachel Kuske, Sam P. Brown

## Abstract

Antibiotics are a cornerstone of modern medicine, targeting pathogen cells by disrupting essential cellular processes. However, standard antibiotic susceptibility metrics (e.g. MIC) and textbook models neglect transient dynamics and density-dependent effects, despite their ubiquity in nature. In clinical infections, where bacterial populations are the units we treat, this can increase the risk of under treatment. To address this gap, we generate high resolution optical density time series data for *Pseudomonas aeruginosa* (3 antibiotics, 12 doses, 7 inoculum sizes, 4x replication), enabling gradient estimation and gradient-based model parameterization. We develop a dynamics-led computational pipeline that (1) evaluates population scale ordinary differential equation models in the context of estimated time derivative data, and (2) classifies transient dynamics in dose-inoculum space using unsupervised clustering. Applied to our data, the pipeline identifies an ordinary differential equation model with a saturating antibiotic-loss term and a threshold-dependent weak Allee term that recapitulates and quantifies classic rate, yield, and inoculum effects of antibiotics. In addition, our model and clustering approach suggest a set of novel metrics, defining thresholds separating distinct dynamical regimes. Beyond antibiotic data sets, our approach utilizing a derivative-based fitting algorithm and clustering of derivative trajectories is applicable to any biological time series with controlled perturbations and variable initial conditions.

**Author summary:** We show that standard math models and antibiotic susceptibility metrics fail to capture the regimes of dynamical behavior that result from combined antibiotic and inoculum effects when *Pseudomonas aeruginosa* (PAO1) is exposed to antibiotics. Using iterations of forward and data-driven modeling, we highlight the importance of transient dynamics, derivative-based model fitting, and higher-order nonlinearities in quantifying bacterial dynamics under perturbation. We identify an ordinary differential equation-based model that describes the observed inoculum effect as a type of weak Allee effect (positive density-dependence) and also captures antibiotic effects on population growth rate and yield governed by a saturating loss function. We show that these results generalize across antibiotic mechanisms of action and highlight the importance of fitting models using dynamics-based algorithms. Finally, we explore a clustering method for bacterial dynamics that, in combination with our mathematical model, advises a set of novel metrics for measuring antibiotic susceptibility.

## Introduction

Antibiotics impact bacterial cellular growth and survival through inhibition of essential cell processes that are unique to bacterial cells. These processes include specific components of cell wall, protein, and DNA synthesis, and essential metabolic pathways [1]. Despite a strong molecular scale understanding of how antibiotics target bacteria, effects on population scale growth—the scale at which we treat—are less well defined [2].

Inoculum effects, or antibiotic induced density dependence, are commonly observed for many combinations of bacteria and antibiotics [3–6], and can be described as an increase in minimum inhibitory concentration (MIC) with an increase in inoculum or initial bacterial population size [7]. In terms of population density, inoculum effects can present as growth bistability, where larger bacterial populations are able to grow at antibiotic concentrations where smaller populations would be inhibited [8–10]. Possible mechanisms for inoculum effects are related to individual and population level behavior [11], including resistant sub-populations or persisters [12–15], titration effects [4, 16–19], degradation [9, 15, 20], and spatial structuring [21]. While there is a lack of consensus as to the underlying mechanisms of inoculum effects, standard antibiotic susceptibility testing protocols are defined based on a standard inoculum size and neglect transient bacterial population dynamics (Fig 1) [16]. Similarly, mathematical models of pharmacodynamics struggle to capture large variation in initial conditions with a single functional form [8, 13, 16].

**Fig 1.**
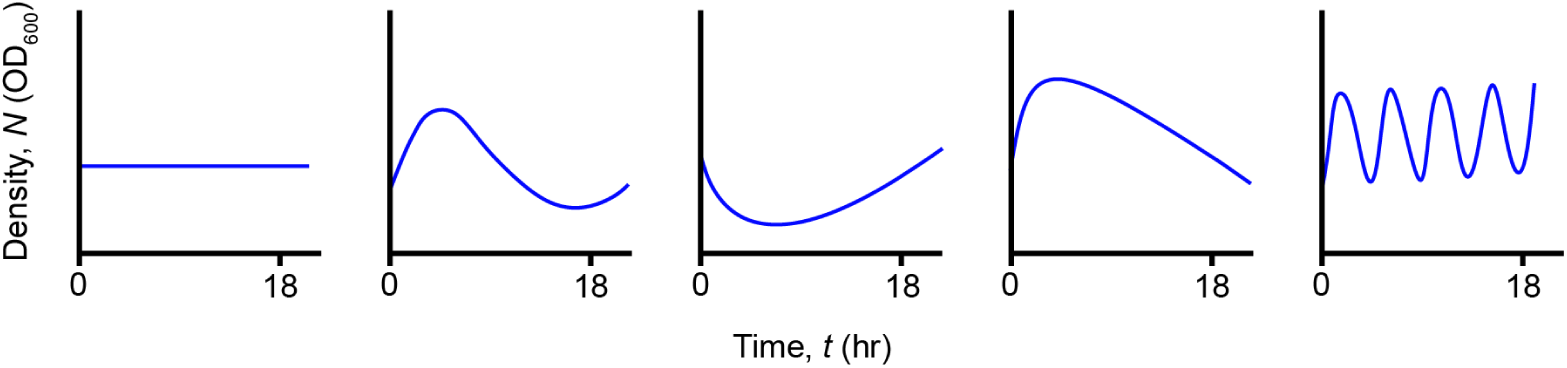
Standard protocols for calculating MIC in practice neglect transient dynamics. Example trajectories of bacterial density under antibiotic exposure vs. time highlight the importance of considering transient dynamics when evaluating bacterial growth. Under standard protocols for measuring MIC [22–25], all five trajectories could be interpreted identically despite showing diverse transient dynamics.

In this study, we investigate the combined effect of antibiotic exposure and inoculum size on bacterial population growth, primarily using the broad spectrum carbapenem antibiotic meropenem, which is commonly used for more serious *Pseudomonas aeruginosa* hospital infections [26, 27]. Using high temporal resolution experimental data of *P. aeruginosa* grown under varied antibiotic dose and inoculum size, we look to answer the following questions: (1) How does inoculum size impact transient bacterial population dynamics under antibiotic exposure? (2) Can we define new metrics for antibiotic susceptibility that better reflect inoculum effects and transient dynamics? and (3) What methods are necessary to capture qualitative and quantitative population-level impacts of perturbations and varied initial conditions? To address these questions, we present and analyze a number of mathematical models from existing theory, and then apply them to data in a sequential fashion with increasing complexity. The goal is to efficiently capture both antibiotic and inoculum effects in our data set while also developing an innovative computational pipeline that integrates familiar models with derivative-based model fitting and unsupervised clustering.

Through iterations of data-driven modeling, we show that standard models fail to capture the regimes of dynamical behavior that result from combined antibiotic and inoculum effects on *P. aeruginosa* and identify a new composite model that defines an antibiotic dose threshold for inoculum effects. Our proposed model characterizes the effect of antibiotics on population density with a saturating loss function and describes the observed inoculum size impacts as a weak Allee effect [28, 29], where the population experiences positive density dependent growth that offsets high antibiotic exposure, capturing modifications to *P. aeruginosa* growth rate and yield due to meropenem exposure. We highlight the importance of transient bacterial population dynamics and present a clustering method for temporal dynamics that also suggests a density dependent measure of antibiotic susceptibility. Finally, we present the generalizability of our results by applying our analyses to the population dynamics of *P. aeruginosa* under the exposure of antibiotics with distinct mechanisms of action, tobramycin and tetracycline. Through this process, we emphasize the importance of careful model selection, consideration of data type and resolution, and incorporation of bacterial population dynamics (e.g. *dN/dt*), not just density measures *N* (*t*), in our inference methods.

## Results

### Antibiotic susceptibility metrics neglect transient dynamics

Current practices for measuring antibiotic susceptibility and predicting bacterial growth dynamics typically rely on the use of MIC and the classic logistic growth model, respectively. Our initial theoretical and experimental investigations reveal that while these classical approaches can capture qualitative declines in both growth rate and growth yield, they neglect more complex transient dynamics that can lead to alternate treatment outcomes.

First, we investigate the microbiological concept of MIC: the lowest concentration of an antibiotic that prevents overnight bacterial growth and can be estimated via standard protocols of antibiotic susceptibility testing [5, 22–24]. Standard protocols for measuring MIC rely on identifying the lowest antibiotic concentration that prevents visible bacterial growth after 16-20 hours [22], when grown using a clinical standard inoculum of 5 × 10^5^ CFU/mL [25, 30] and specific growth environment.

Clinically, MIC provides a strategy for classifying a population as ‘susceptible’ or ‘resistant’ based on whether the MIC is less than or greater than the MIC breakpoint established by CLSI [31]. However, reliance on MIC for determining susceptibility of a population has its shortcomings. Human bacterial infections are dynamic and complex, whereas the protocol for determining MIC relies on a standard inoculum concentration and growth environment that is unlikely to reflect infection conditions. Measuring MIC at higher inocula is problematic, particularly if the initial density is greater than the threshold for visible growth. In addition to inoculum effects, MIC measurements are also potentially sensitive to the time window chosen. Fig 1 provides a simple schematic where we show five highly distinct dynamical scenarios that would all lead to the same MIC estimate using standard methods. We hypothesize that transient dynamics are instrumental in understanding how effective the bactericidal or bacteriostatic effect of an antibiotic is on a population.

To assess the importance of transient dynamics given antibiotic perturbation, we first introduce the classic logistic growth model, commonly used to model microbial population growth. We define the ordinary differential equation (ODE),

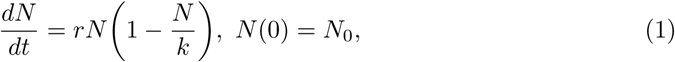

where *N* (*t*) is bacterial density at time *t*, *r* is the maximal growth rate, and *k* is the carrying capacity given environmental conditions (Table 1). More generally, we interpret the carrying capacity as the population “growth yield”, a term we use in the following to describe the bacterial density as *t* → ∞. In Eq 1, the growth yield is simply the equilibrium density, *N* ^∗^ = *k* (Table 1, Section 1.1 in S1 Text). We modify Eq 1 to include antibiotic perturbation in the form of an additional loss term [12, 16, 32],

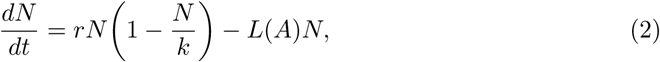

**Table 1.** Model parameters and terminology.

| Parameter | Description | Units |
| --- | --- | --- |
| $N(t)$ | bacterial density at time $t$ (hr) | OD<sub>600</sub> (density) |
| $\Delta t$ | time step for numerical simulation | hr |
| $N_0$ | initial bacterial density or inoculum concentration | OD<sub>600</sub> (density) |
| $A$ | antibiotic concentration | μg/mL |
| $r$ | maximal growth rate | hr<sup>-1</sup> |
| $k$ | carrying capacity | OD<sub>600</sub> (density) |
| $N^*$ | equilibrium density or “growth yield” | OD<sub>600</sub> (density) |
| $r(A)$ | effective growth rate | hr<sup>-1</sup> |
| $k(A)$ | effective carrying capacity (approx. $N^*$) | OD<sub>600</sub> (density) |
| $L(A)$ | effective antibiotic loss rate | hr<sup>-1</sup> |
| $l$ | antibiotic loss coefficient | hr<sup>-1</sup> |
| $A_{MIC}$ | MIC | μg/mL |
| $h$ | Hill coefficient | — |
| $A_{50}$ | half maximal effect concentration | μg/mL |
| $a$ | Allee threshold | OD<sub>600</sub> (density) |

where *L*(*A*) is a function describing the rate of population decline due to antibiotic concentration, *A*. For simplicity, we initially consider a simple linear rate in line with the concept of MIC,

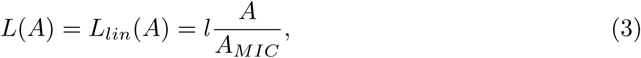

where the loss coefficient is defined as *l* = *r* such that the effective growth rate will go to 0 as *A* → *A_MIC_*.

Simulating bacterial population density under varied antibiotic concentration using Eq 2, we see that both the growth rate in the exponential growth phase, and the growth yield of the population decline with increasing antibiotic concentration (Fig 2A).

**Fig 2.**
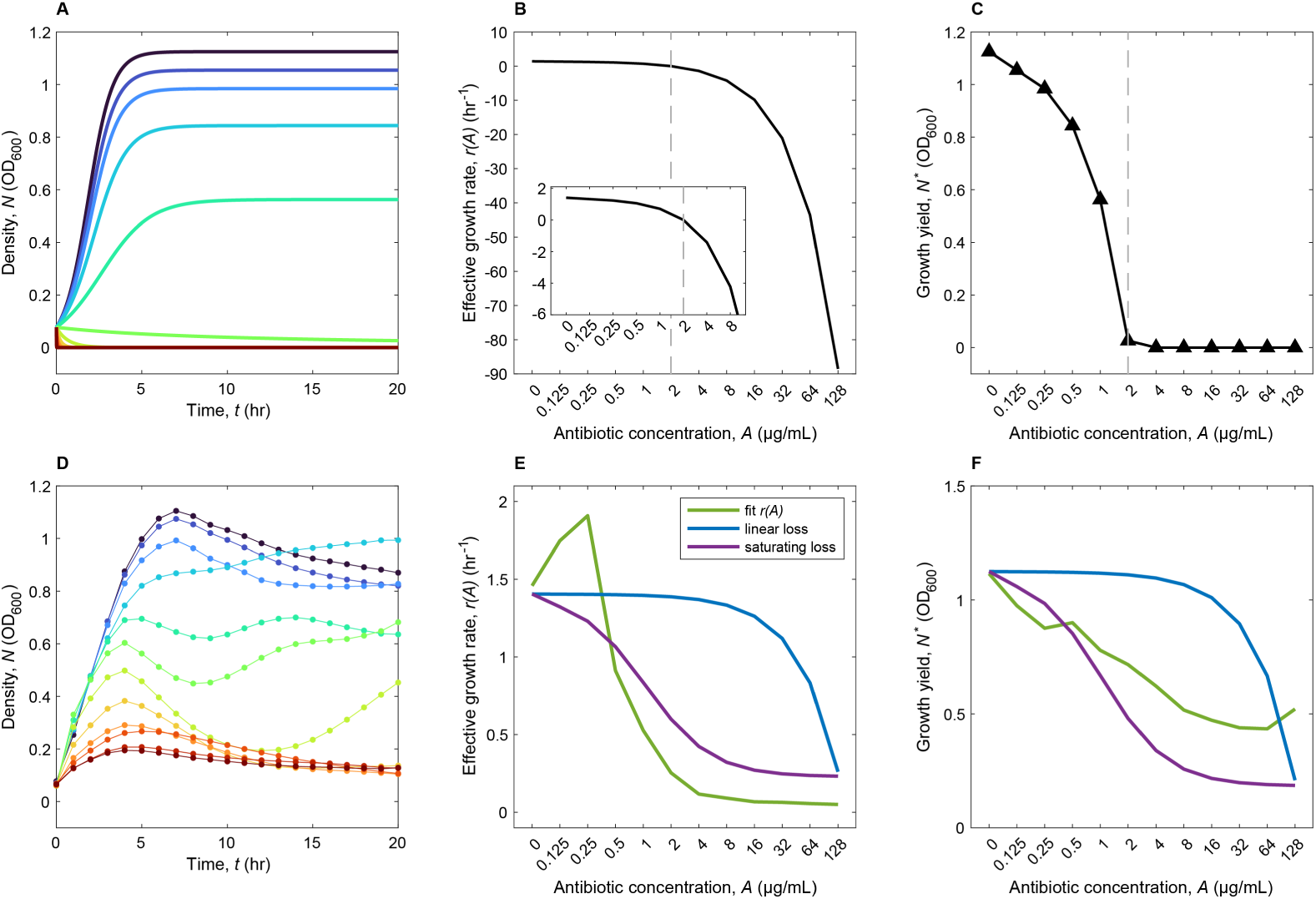
Effects of antibiotic perturbation on bacterial growth rate and growth yield. A) Simulated density data for Eq 2 given varied antibiotic concentration *A*. Colored lines represent different antibiotic concentrations with dark blue representing no antibiotic and dark red showing maximum antibiotic. (B) Simulated effective growth rate *r*(*A*) vs. antibiotic concentration *A* for Eq 2. (C) Simulated growth yield *N* ^∗^ vs. antibiotic concentration *A* for Eq 2. The gray dashed line in panels B and C, shows the value of MIC, *A_MIC_* = 2 µg*/*mL, where *r*(*A*) = 0, used for model simulation. Panels D-F compare these simulated predictions to data. (D) Experimental data for bacterial density vs. time for the inoculum size and range of meropenem exposure corresponding to the simulated data in panel A. Panels E and F show the effective growth rate and growth yield, respectively, vs. meropenem concentration, from model fitting for (i) an effective growth rate and growth yield for each antibiotic concentration separately (green, Eq 1 with Algorithm 1), (ii) a linear antibiotic effect (blue, Eq 2 with *L*(*A*) defined by Eq 3, fit with Algorithm 2), and (iii) a nonlinear (saturating) antibiotic effect (purple, Eq 2 with *L*(*A*) defined by Eq 4, fit with Algorithm 2). Simulation parameters for panels A-C: *r* = 1.4052, *k* = 1.1249, *l* = 0.7026, *A_MIC_* = 2, *A* = [0, 0.125, 0.25, 0.5, 1, 2, 4, 8, 16, 32, 64, 128], *N*_0_ = 0.0771, Δ*t* = 0.01. Fit parameters for panels E-F: (green) as shown; (blue): *r* = 1.4052, *k* = 1.1249, *l* = 0.0726, *A_MIC_* = 8.1194; (purple): *r* = 1.4052, *k* = 1.1249, *l* = 1.1764, *h* = 1.2117, *A*_50_ = 1.0537.

Mathematically, declines in growth rate are captured in Eq 2 by the function *r*(*A*) = *r* − *L*(*A*), shown in Fig 2B. In Fig 2C, declines to growth yield are captured by the modification of the equilibrium point by the antibiotic, 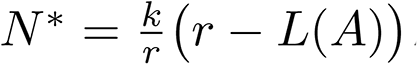. The gray dashed line in Fig 2B and 2C corresponds to the MIC value defined for simulation of Eq 2, *A_MIC_* = 2 µg*/*mL, indicating the transition from positive to negative effective growth rate and the lowest dose with zero growth yield as *t* → ∞. Similarly, the time series trajectories for antibiotic exposures at MIC and higher (bright green to dark red, where density *N* → 0 as *t* → 20) in Fig 2A show no increase in optical density (OD) over 16-20 hours, consistent with standard protocols.

In order to assess whether Eq 2 realistically captures bacterial population dynamics, we consult our experimental data tracking the density of bacterial pathogen, *P. aeruginosa* via hourly measures of OD, under 12 different doses of meropenem for 7 inoculum concentrations, each in 4-fold replication. Detailed methods are outlined in Materials and methods, and time series data is shown in S1 Fig. In Fig 2D, we show the time series trajectories for a single inoculum concentration under the same levels of meropenem exposure used for our model predictions in Fig 2A.

While we observe declines in growth rate and growth yield in Fig 2D similar to our model predictions in Fig 2A, the temporal dynamics are much more complex, exhibiting trajectories that more closely resemble scenarios from Fig 1 and fail to be eliminated at the highest antibiotic dose. This suggests that while the overall functional form of Eq 2 is able to capture qualitative declines in both growth rate and yield observed in the temporal dynamics (Fig 2A vs. Fig 2D) in contrast to other models which capture declines in growth rate only [4, 33, 34], the effect of antibiotic exposure is overestimated and likely, oversimplified in the model, particularly at moderate levels of antibiotic exposure.

### Meropenem exposure nonlinearly impacts *P. aeruginosa* growth rate and yield

Having sketched the broad qualitative expectations of the effects of antibiotic exposure on bacterial density, we next consider the functional form of antibiotic effects. By utilizing theory and experimental data, we seek functional forms that represent the underlying biological mechanisms driving population dynamics, while still obtaining interpretable and generalizable models. First, we focus solely on the effects of antibiotic concentration, initially assuming that differences in inoculum present in our data set represent natural variation in initial condition, without explicitly impacting dynamics (Algorithm 1, Materials and methods and Section 2.2 in S1 Text).

As we noted in the previous section, inspection of the time series plots (Fig 2D, S1 Fig) suggests that both growth rate and growth yield are declining with increasing antibiotic concentration, but that a simple linear loss term using MIC (Eq 3) may be insufficient. To investigate this qualitative assessment, we map the empirical response of effective growth rate, *r*(*A*) and growth yield, *N* ^∗^, to increasing antibiotic concentration by producing separate fits for each antibiotic concentration. Here, we use the fitting protocol described in Algorithm 1 (see Section 2.2 in S1 Text), which assumes the dynamics of *N* (*t*) are the same regardless of inoculum concentration *N*_0_ and yields an effective growth rate *r*(*A*) and effective carrying capacity *k*(*A*), which approximates the growth yield *N* ^∗^, for each treatment (green lines, Fig 2E-F, Table 1). We see that both effective growth rate and growth yield decline with increasing antibiotic concentration (*dr*(*A*)*/dA <* 0, *dk*(*A*)*/dA <* 0). Biologically, this makes sense as beyond altering growth rate, antibiotics have been shown to impact growth efficiency and yield [5, 13, 35], meaning that not only will the population grow more slowly given exposure, but it won’t be able to maximize its density to the carrying capacity in the absence of antibiotic exposure, *k*.

Next, we compare these results to two functional forms for the loss rate with respect to antibiotic concentration *L*(*A*) in Eq 2, both of which also exhibit declines in effective growth rate and yield. One is the linear form using MIC introduced in the previous section (Eq 3) and the other is a nonlinear, saturating loss function using half maximal concentration, *A*_50_. In the second, we define the saturating antibiotic loss rate as,

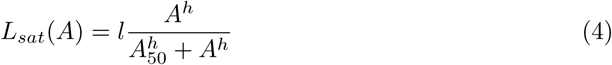

where the loss coefficient *l* now describes the maximal loss rate due to antibiotic and *h* is the Hill coefficient. We note that both forms are commonly used in the microbial and pharmacodynamic literature, with the linear form commonly being used to approximate the nonlinear, saturating function form [36, 37]. More complex saturating forms exist utilizing both MIC and *A*_50_ [4, 5, 16], but these can be simplified to Eq 4 and give the same dynamics just with a different combination of parameters [16, 33].

We fit Eq 2 with the linear (Eq 3) and saturating (Eq 4) loss rates using the fitting protocol defined in Algorithm 2, which looks to fit a common parameter set to all of the dynamics (approximate derivatives *dN/dt*) of our data where antibiotic concentration *A* is treated as an independent variable (Materials and methods and Section 2.3 in S1 Text). Then, we compare the effective growth rate *r*(*A*) = *r* − *L*(*A*) (Fig 2E) and the growth yield *N* ^∗^ (Fig 2F) with the results from fitting the data using Algorithm 1.

We note that using Algorithm 2 allows us to more efficiently capture the dynamics of the entire data set by using fewer parameters. For example, we describe the data with 4 parameters for Eq 2 with *L_lin_*(*A*) and 5 parameters for Eq 2 with *L_sat_*(*A*) versus 24 parameters when we fit individual *r*(*A*) and *k*(*A*) parameters for each antibiotic dose *A* (green curves in Fig 2E-F). Additionally, Algorithm 2 estimates parameters with “gradient matching” [37], attenuating the effects on the minimization problem from variation of the initial densities over 3 orders of magnitude, by instead focusing on the rate of change in density.

Fig 2E shows that in both the case of linear loss (blue line, Eq 2 with Eq 3) and saturating loss (purple line, Eq 2 with Eq 4), we roughly approximate the behavior of the fit effective growth rate (green line) at low antibiotic concentrations, and also offer similar predictions at the highest antibiotic concentration tested. However, the linear case largely underestimates the effect of increasing antibiotic concentration until much higher concentrations are reached, whereas the saturating loss only mildly underestimates the effect at intermediate values (Fig 2E). Given that we describe the data set with fewer parameters, we expect declines in quantitative agreement. However, the saturating loss form generally captures the concave up shape of the fit effective growth rate curve that the linear loss form misses.

In Fig 2F, we consider the change in growth yield, 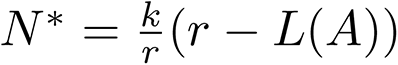 (Table 1) vs. *A* as predicted by Eq 2 with saturating antibiotic loss (Eq 4, purple line). We see that it approximately captures the behavior of the fit effective carrying capacity *k*(*A*) (green line), although it tends to overestimate the effect of *A* on *k* as antibiotic concentration increases. Similar to the results for the effective growth rate, tracking growth yield versus antibiotic concentration for the model with linear antibiotic loss (blue line, Eq 2 with *L_lin_*(*A*)) underestimates the effect of antibiotic on the fit effective carrying capacity (green line, Eq 2F).

While linear approximations of antibiotic loss may be adequate in some cases, for example at very low or very high *A*, the effect of antibiotic exposure on bacterial population dynamics with respect to growth rate and yield is more accurately described by the saturating loss function (Eq 4). For clarity, we combine Eq 2 and Eq 4,

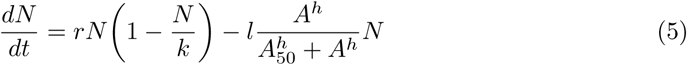

and refer to this model as “logistic growth with saturating loss”. Next, we use this model as our baseline to relax the assumption that inoculum size has no explicit effect on dynamics in conjunction with antibiotic exposure.

### The need for minimal models that capture inoculum effects

In the previous sections and in the general modeling literature [38–42], we treat differences in inoculum as natural variation in initial density that doesn’t explicitly impact dynamics or parameter values. This is consistent with utilizing a standard inoculum size for antibiotic susceptibility testing: it assumes initial density does not alter outcomes of antibiotic exposure. Yet it is widely recognized that antibiotics can be less effective when used to treat higher densities—a form of positive density dependent growth that is described by microbiologists as an “inoculum effect” [7]. To address inoculum effects, we first ask: what effect does inoculum size, defined here as initial density *N*_0_, have on the dynamics defined by Eq 5?

In Fig 3, we compare simulations of Eq 5 with experimental data under no (*A* = 0 µg*/*mL, black) and intermediate levels of antibiotic exposure (*A* = 2 µg*/*mL, red). In the model simulations (Fig 3A-D), antibiotic exposure alters both the growth rate and growth yield of the population, but inoculum size only alters the time it takes for the population to reach the growth yield regardless of antibiotic dose (see Section 1.1 in S1 Text). (We simulate two additional antibiotic conditions with varied inoculum sizes in S2 Fig, showing that neither the effective growth rate *r*(*A*) = *r* − *L*(*A*) nor growth yield *N* ^∗^ vary with inoculum size at low and high antibiotic exposure.) In the no antibiotic case, the experimental data (Fig 3E-H), shows dynamic similarity across inocula. Visually, the variation with inocula across the experiments with no antibiotic can be approximated by the simulations in Fig 3A-D, capturing the reduced time to reach maximal growth yield, while the the maximal growth rate and yield appear approximately constant regardless of *N*_0_.

**Fig 3.**
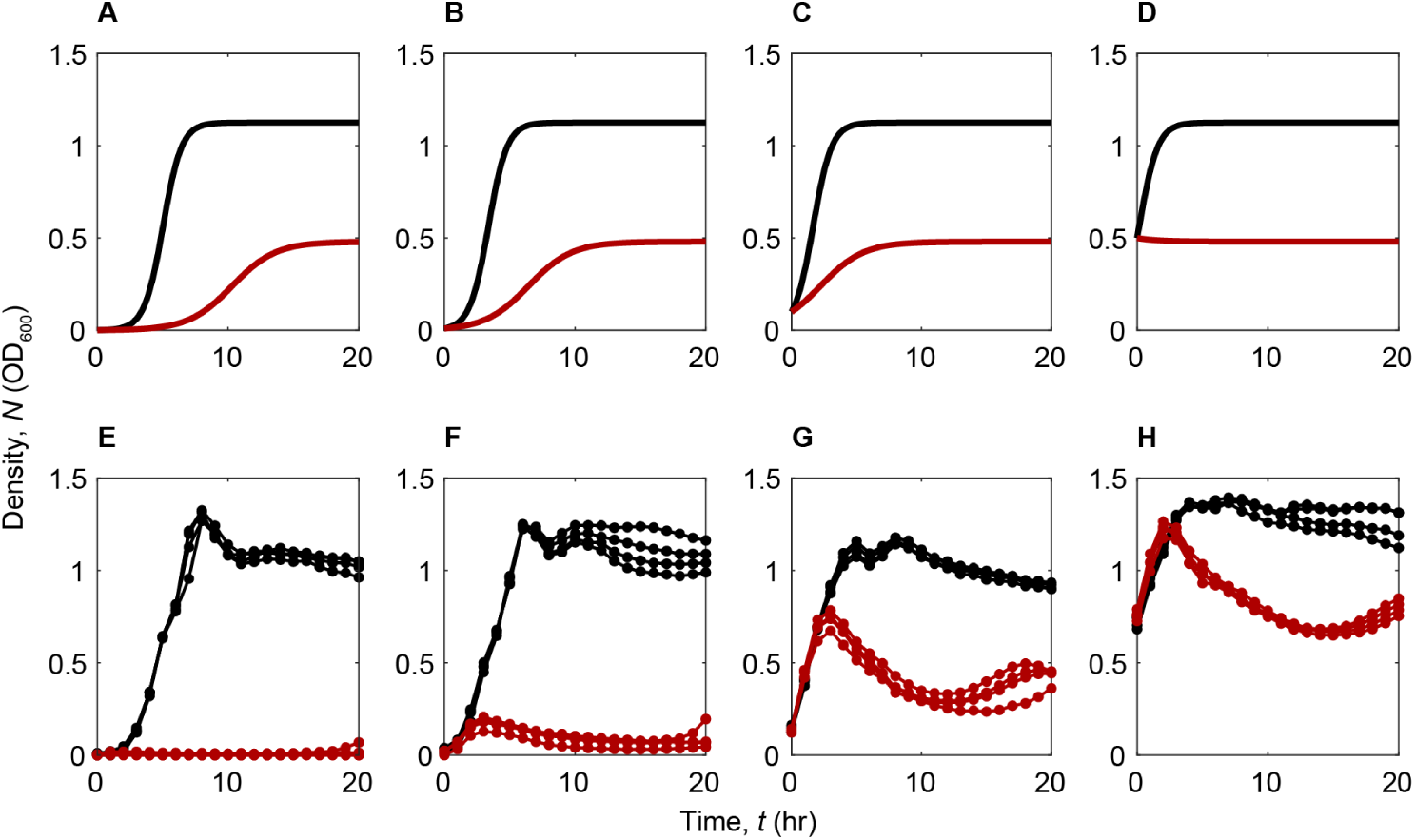
Simulation of Eq 5 in the presence and absence of antibiotic exposure compared with experimental data. Black lines show the no antibiotic case (*A* = 0 µg*/*mL) and red lines indicate intermediate antibiotic exposure (*A* = 2 µg*/*mL). Inoculum size increases identically from Panel A to D and Panel E to H where *N*_0_ = 0.001, 0.01, 0.1, 0.5. Panels A-D: simulated data from Eq 5 shows no impact of variation in inoculum size on growth yield. Panels E-H: corresponding experimental data shows clear variation in growth rate and yield under different inoculum conditions. Parameters for simulated data: *r* = 1.4052, *k* = 1.1249, *l* = 1.1764, *h* = 1.2117, *A*_50_ = 1.0537, *A* = 0, 2, Δ*t* = 0.01.

We contrast these observations with the experimental data under moderate antibiotic exposure (*A* = 2 µg*/*mL), where the dynamics for different inocula vary substantially with respect to growth rate and growth yield (red lines in Fig 3E-H). Given antibiotic exposure, the lowest inoculum size has little to no population growth over 20 hours, remaining at very low densities (Fig 3E). In contrast, Fig 3F-H show that increased inoculum size leads to elevated final densities that are greater than or equal to *N*_0_. As in the no antibiotic case, the length of time to maximal growth yield predictably decreases with increasing inoculum size, as the population starts closer to *N* ^∗^. S3 Fig provides examples of similar behavior under other levels of antibiotic exposure.

Comparing the experimental data for moderate antibiotic exposure with results from Eq 5 in Fig 3A-D (red lines) indicates that the model is not capturing any sort of dynamical inoculum effect observed experimentally.

### Variation in dynamics stems from combined antibiotic and inoculum effects

Fig 3 indicates that the effect of meropenem on *P. aeruginosa* is modified by inoculum size, consistent with the inoculum effect literature [3, 5, 6, 8–10]. To provide a quantitative test for inoculum effects, we use unsupervised learning techniques (*k*-means [43]) to cluster all time series (12 antibiotic concentrations *A* × 7 inoculum doses *N*_0_) into similar groups based on their dynamics (*dN/dt*). If inoculum size has no effect, we expect the time series to cluster according to levels of *A* only, while antibiotic-dependent inoculum effects will result in clustering by both *A* and *N*_0_.

Fig 4 shows the results of applying *k*-means clustering to the approximate derivative trajectories *dN/dt* from our data set for each antibiotic, inoculum treatment. We see that the dynamics can be optimally separated into 5 regimes (Fig 4A, see Materials and methods for clustering details). Further, we can relate the corresponding families of trajectories (Fig 4B-K) to biological interpretations. Trajectories in Cluster 1 (light blue) correspond to no to low growth conditions, where sufficiently high antibiotic exposure for a given inoculum leads to reduced growth rate and yield (light blue in Fig 4A, B, and G). For the lowest inocula tested, the boundary of Clusters 1 and 3 approximately agrees with the MIC from antibiotic susceptibility testing, as indicated by the white dashed line labeled MIC = 2 µg*/*mL. However, the boundary between Cluster 1 and the other clusters diverges from MIC as the inoculum size is increased, flagging the presence of an inoculum effect. Specifically, we see that progressively higher antibiotic concentrations are required to produce pathogen dynamics in the Cluster 1 regime of full control (*dN/dt* persistently close to zero). We also note that for inocula with OD above 0.01, the antibiotic concentration required for control as in Cluster 1 is no longer achievable, as it exceeds the permitted safe dose as defined by CLSI [31] (white dashed line labeled at *R* ≥ 8 µg*/*mL, Fig 4A).

**Fig 4.**
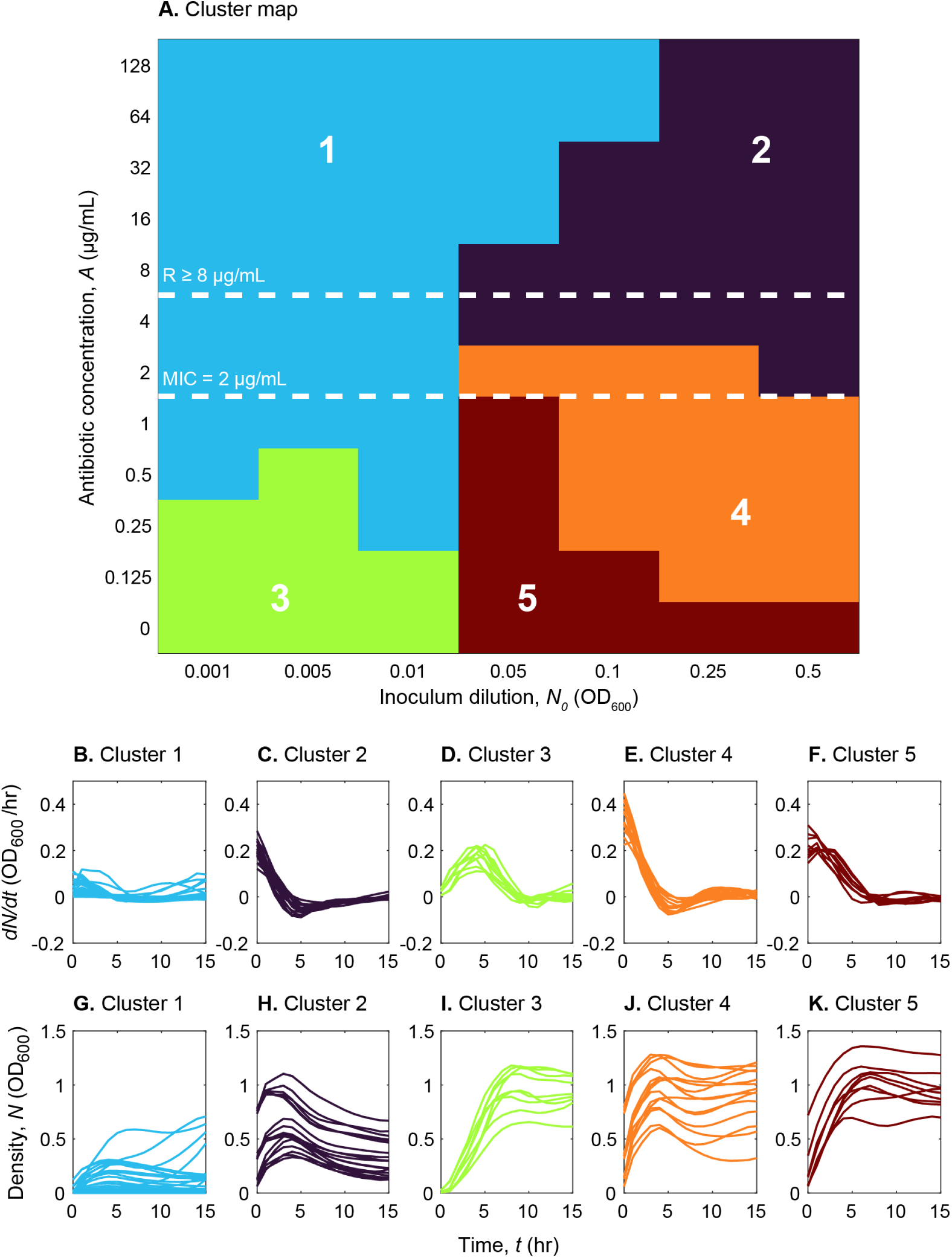
Curves clustered using *k*-means show 5 regimes of dynamic behavior as decided by combinations of both *A* and *N*_0_. Colors and labels denote clusters in antibiotic, inoculum treatment space. Panel A shows treatments in the space of antibiotic vs. inoculum, colored by cluster number. White dashed lines denote MIC determined via standard antibiotic susceptibility testing, labeled MIC = 2 µg*/*mL (Materials and methods) and the CLSI breakpoint for resistance of *P. aeruginosa* to meropenem, labeled *R* ≥ 8 µg*/*mL [31]. Panels B-K show the corresponding time series for density *N* (*t*) and approximate derivatives *dN/dt* for each cluster, depicting that trajectories in Cluster 1 exhibit no to low growth dynamics and remaining clusters depict intermediate and near normal growth.

The other 4 clusters span intermediate to near normal growth (sub-MIC growth dynamics), showing that higher initial density permits near normal growth at higher antibiotic concentrations (Fig 4A, C-F, H-K). While we can combine these groups as one “near normal” growth regime, using the optimal number of clusters highlights that sub-MIC population dynamics are highly variable and clearly density-dependent (Fig 4C-F, H-K). The core results are conserved when we cluster on the density curves *N* (*t*) (S4 Fig). There is again a clear diagonal boundary between “growth” and “no growth” in antibiotic vs. inoculum space. We interpret this boundary as akin to a density dependent MIC characteristic of inoculum effects: MIC increases with increasing inoculum. However, for clusters based on *N* (*t*), the initial and final densities dominate over transient dynamics, leading to lower resolution of the dynamical regimes (2 clusters in (S4 Fig) vs. 5 in Fig 4) and a boundary between Clusters 1 and 2 that underestimates the boundary between Cluster 1 and all others observed in Fig 4A. By clustering on *dN/dt* curves instead of *N* (*t*), we capture the higher doses needed to obtain Cluster 1 control of pathogen growth at intermediate to high inoculum size that would be otherwise underestimated using *N* (*t*).

### Combined antibiotic and inoculum effects on *P. aeruginosa* density can be captured with a weak Allee effect

Having shown that the effect of meropenem on *P. aeruginosa* is modified by inoculum size, consistent with the commonly described inoculum effect, we turn to the question: what are the functional forms that best capture the mechanism of inoculum effects?

First, we hypothesize that inoculum effects are the result of positive density dependence or the inoculum concentration shifting the system into regimes with different dynamics or alternative equilibria, as supported by the evidence of distinct clusters in Fig 4.

To obtain our candidate models, we consider several modifications of Eq 5, each incorporating a different form of density dependence that may capture the underlying mechanisms of inoculum effects, rather than introducing inoculum as an independent variable [16]. Our candidate models describe modifications to growth rate, growth yield, and antibiotic dependent loss via biological processes that reduce antibiotic effects as density increases. Section 1.2 in S1 Text describes the modifications of Eq 5 and the biological and mathematical mechanisms that make them relevant for describing inoculum effects. While the predicted behavior of these models may not completely quantify the observed dynamical behavior in our data set, we focus on identifying aspects of our models that efficiently capture the key elements of the experimental population dynamics. In turn, this will allow us to map the biological mechanisms represented by the functional forms to hypotheses about the influence of the inoculum.

We fit the candidate models using the fitting protocol described in Algorithm 2 (Section 2.3 in S1 Text). By fitting our models via dynamics versus densities, we are able to obtain a single parameter set for each model over all data, yielding a functional form that describes the impacts of both antibiotic concentration and inoculum density. Here, the critical benefit of taking a dynamics-based fitting approach is that it allows us to capture signatures of growth dynamics at transient density levels. As indicated by the clusters in Fig 4, these transients are more effective at identifying differences due to inocula given the large variation in initial and long time density levels. Because Algorithm 2 is based on fitting a single functional form across all treatments, we obtain a much smaller number of parameters than when we parameterize models via Algorithms 1 or 3 (Sections 2.2 and 2.4 in S1 Text), which fit each antibiotic level or treatment separately. Our model prediction error will be predictably higher using Algorithm 2, but we can ensure that our data has sufficient resolution for model parameterization and that our model is generalizable and biologically interpretable. In contrast, with Algorithms 1 or 3, we could fit a model perfectly with an increasing number of parameters but this does not typically enhance our understanding of the rules governing system behavior.

We compare the candidate model predictions with experimental data and the logistic growth model with saturating loss in (Eq 5) by calculating the root mean squared error (RMSE) for each treatment,

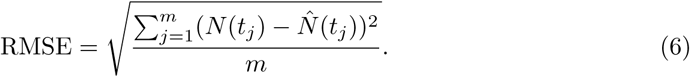

Specifically, RMSE is obtained by comparing the model simulation using fit parameters at time *t_j_* (*N*(*t_j_*)) with the corresponding data *N* (*t_j_*) summed across time points, for each inoculum and antibiotic combination, for each model tested (Section 4 in S1 Text).

While we parameterize our models using the derivatives instead of densities, we use Eq 6 to evaluate the model results with respect to our experimental data to provide a standard comparison: do model dynamics produce the variability in density measures we observe?

Looking at Fig 5A, we see that while the logistic growth model with saturating loss approximately captures bacterial density across most of the treatment space, it struggles to reproduce growth rate and yield observed experimentally at inocula extremes. In contrast, our candidate models all perform better at high antibiotic exposures than Eq 5 (Fig 5B, S7 Fig), largely due to their ability to more closely approximate less severe declines in growth yield. This supports the existence of positive density dependencies that offset strong antibiotic effects.

**Fig 5.**
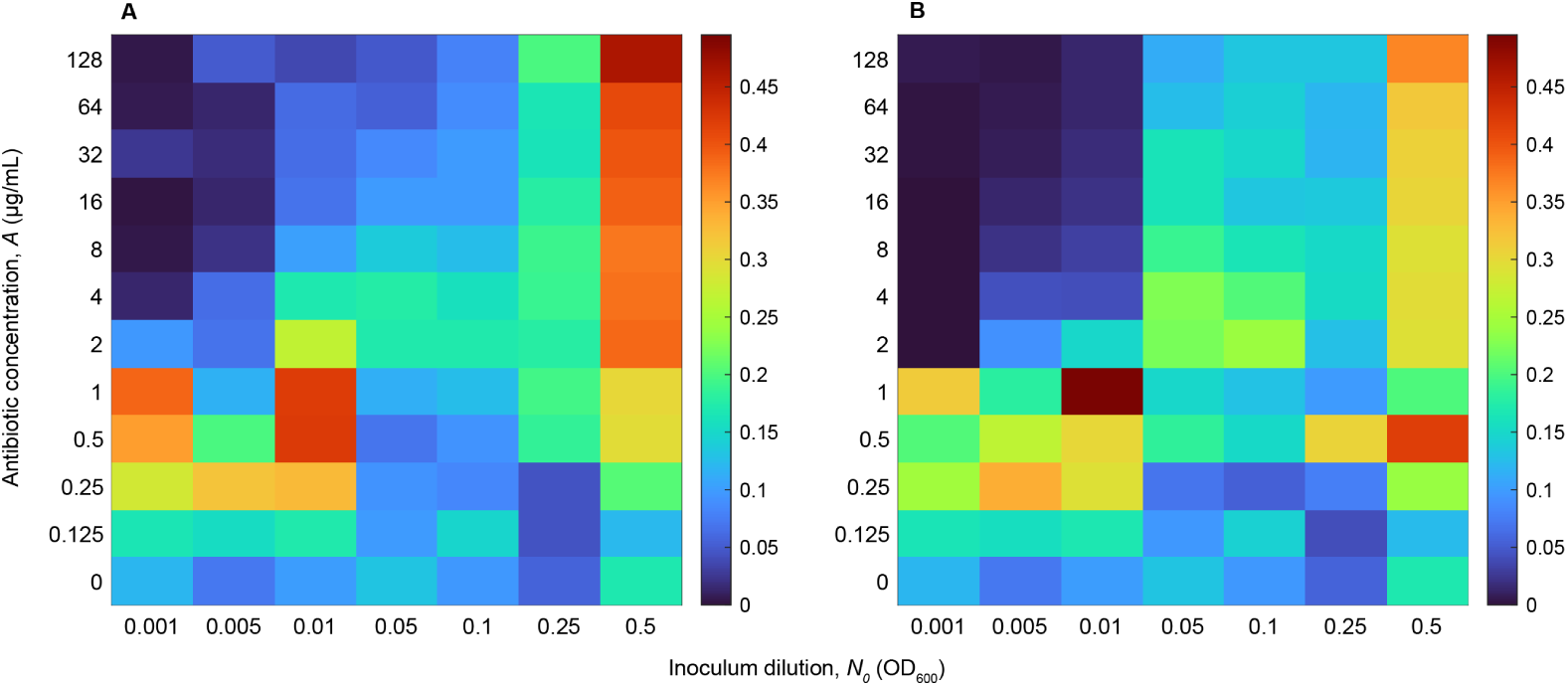
Heat maps of RMSE for data from all treatments (antibiotic concentration and inoculum size combination). Panels A and B compare the RMSE across treatment space of the prediction from the logistic growth model with saturating antibiotic loss (Eq 5, Panel A) and the antibiotic threshold dependent weak Allee effect model (Eq 7, Panel B) fit via Algorithm 2. Parameters for Panel A follow from Fig 2. Parameters for Panel B are defined in Table 2 for meropenem. S5 Fig and S6 Fig show the corresponding time series trajectories for each model vs. data. S7 Fig and S8 Fig show the RMSE and select time series trajectories for the additional candidate models.

**Table 2.** Model parameters values from fitting the antibiotic threshold dependent weak Allee effect model (Eq 7) for meropenem, tobramycin, and tetracycline data sets using Algorithm 3 for parameters *r* and *k*, and Algorithm 2 for all other parameters.

| Antibiotic | $r$ ($\text{hr}^{-1}$) | $k$ ($\text{OD}_{600}$) | $l$ ($\text{hr}^{-1}$) | $h$ | $A_{50}$ ($\mu\text{g/mL}$) | $a$ | $A_{thresh}$ ($\mu\text{g/mL}$) |
| --- | --- | --- | --- | --- | --- | --- | --- |
| Meropenem | 1.4052 | 1.1249 | 1.4924 | 1.5302 | 0.7751 | 0.7161 | 1 |
| Tobramycin | 0.7288 | 1.1976 | 0.7576 | 1.5657 | 0.7111 | 0.6557 | 2 |
| Tetracycline | 0.7288 | 1.1976 | 0.7477 | 1.4446 | 6.2201 | 0.7562 | 8 |

Consistent with our observations in Fig 4, the error maps suggest that the antibiotic dose vs. inoculum space can be divided into different regimes: those where all or most models capture the dynamics and those where certain models outperform others (Fig 5, S7 Fig). This is intuitive given the diversity of dynamics we observe across treatments—it’s difficult for one model to capture both signatures of sub- and super-MIC growth inhibition at once, that is, near normal growth and no to low growth, respectively. Comparing the data to the density predicted by the models, we see that the models almost always fail to capture transient dynamics regardless of treatment condition (S5 Fig and S8 Fig). Specifically, we observe very limited modification of growth rate and yield across inoculum. The model failure is most pronounced around the the diagonal of increasing inoculum size and increasing antibiotic concentration.

By utilizing a “switch” dependent on antibiotic dose, we can provide a composite model for capturing near normal growth dynamics as well as positive density dependence under higher antibiotic exposures. We introduce an antibiotic threshold dependent weak Allee effect into the model, defined as

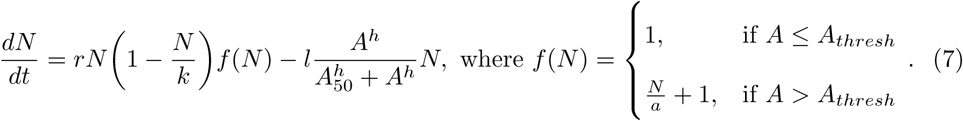

where *f* (*N*) describes the weak Allee effect [29, 44], *a* is the Allee threshold, and *A_thresh_*is the antibiotic threshold governing the “switch”. This model captures both the success of the logistic growth model with saturating loss at lower antibiotic exposures (near normal growth regimes), and the positive density dependence of the inoculum effect at intermediate and high antibiotic exposures (Fig 5B, intermediate to no growth regimes). This success is apparent both in the highest inoculum columns of Fig 5 and in comparisons of the transient dynamics of the logistic growth with saturating loss and weak Allee effect models versus our experimental data (S5 Fig and S6 Fig).

Fig 6 provides additional insight on the contrast between models using the predicted per-capita net growth *g*(*N*),

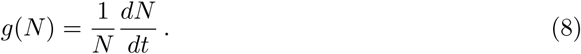

**Fig 6.**
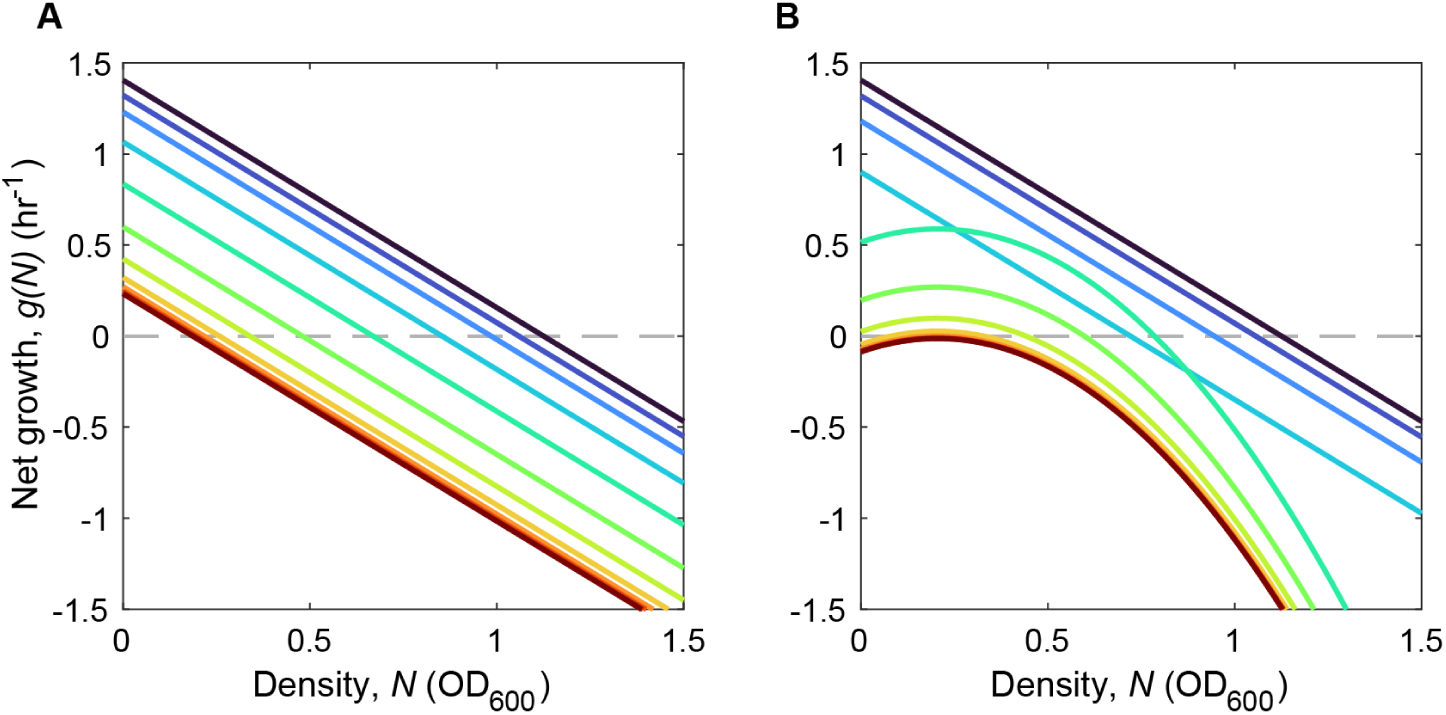
Comparison of per-capita net growth rate for the logistic growth model with saturating loss (Eq 5) and the weak Allee effect model (Eq 7). Net growth vs. density for Eq 5 (panel A) and Eq 7 (panel B) under varied antibiotic exposures (dark blue = lowest antibiotic concentration, dark red = highest antibiotic concentration, *A* = [0, 0.125, 0.25, 0.5, 1, 2, 4, 8, 16, 32, 64, 128]). Gray dashed line denotes *g*(*N*) = 0. Parameter values for qualitative predictions are from model fitting to the full data set via Algorithm 2, see Fig 2 for Eq 5 parameters and Table 2 for Eq 7 parameters.

Specifically, the logistic growth model with saturating loss (Eq 5) shows a linear decrease of *g*(*N*) with density *N* for all antibiotic levels, while the antibiotic threshold dependent weak Allee effect model (Eq 7) shows a nonlinear dependence for antibiotic levels above a threshold, *A > A_thresh_* (Fig 6B), which is fit based on our data as *A_thresh_* = 1 µg*/*mL for meropenem. This nonlinear behavior of (Eq 7) captures the experimental results shown in Fig 3E-H. In particular, over the range *N* ∈ [0, 0.5] of inoculum concentrations, when *A > A_thresh_*, the per-capita net growth rate *g*(*N*) experiences little to no change with increasing density, in contrast to *g*(*N*) for larger values of *N* or when *A* ≤ *A_thresh_*.

### Application to other antibiotics

Up to this point, our analysis has focused on *P. aeruginosa* exposure to meropenem, a bactericidal antibiotic that inhibits cell wall synthesis leading to cell death [27]. While we select meropenem as our focal drug for its relevance in treating severe and often high bacterial load *P. aeruginosa* infections, we ask: are positive density dependent effects observed under meropenem exposure generalizable to antibiotics with other mechanisms of action?

Applying our candidate models (Section 1.2 in S1 Text) to our tobramycin and tetracycline data sets (Materials and methods), we again find that introducing an antibiotic threshold dependent weak Allee effect is able to capture combined antibiotic and inoculum effects across the treatment space (Fig 7). Looking at both the time series trajectories and the RMSE in Fig 7, we see that despite different antibiotic mechanisms of action, *P. aeruginosa* continues to exhibit logistic growth dynamics at low antibiotic concentrations and positive density dependence in the form of a weak Allee effect at higher antibiotic doses. Consistent with meropenem (Fig 5B), we see that the weak Allee model produces low RMSE at the highest inoculum sizes and at high antibiotic concentrations for both drugs (Fig 7).

**Fig 7.**
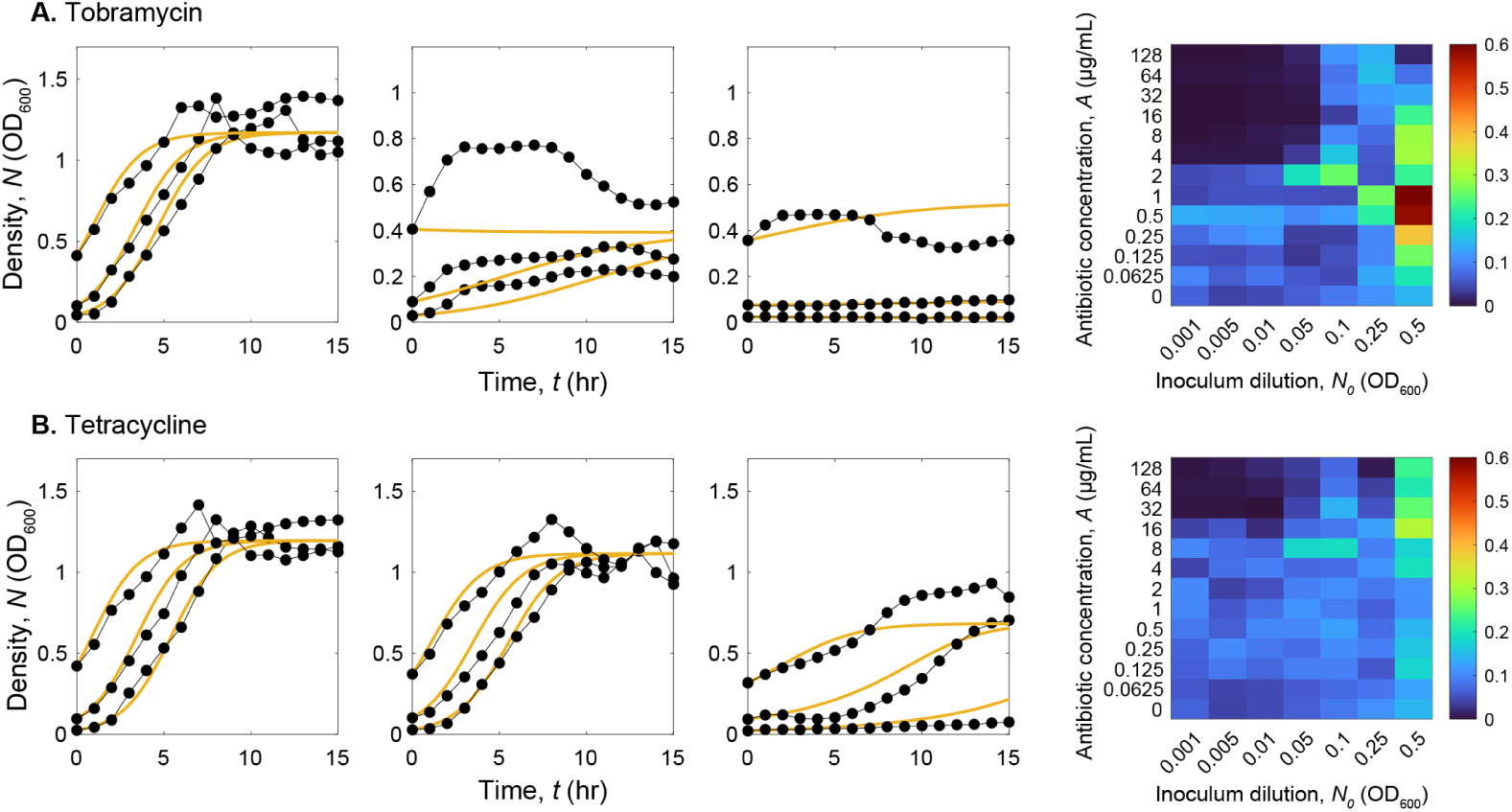
Antibiotic threshold dependent weak Allee effect model captures bacterial dynamics under tobramycin and tetracycline exposures. Time series and corresponding RMSE heatmaps for the antibiotic threshold dependent weak Allee effect model (Eq 7) for *P. aeruginosa* exposed to tobramycin (Panel A) and tetracycline (Panel B). Model prediction in yellow, with model parameters given in Table 2 and simulation time step defined Δ*t* = 0.1, compared to black dotted lines showing experimental data. Antibiotic concentrations: 0.0625, 1, 16 µg*/*mL (time vs. density axes by row, left to right). Inoculum dilutions: 0.005, 0.05, 0.25. Heat maps (far right) display the RMSE (Eq 6) across the space of antibiotic concentration vs. inoculum size for the weak Allee effect model (Eq 7).

We also see similar results when clustering the dynamics of our tobramycin and tetracycline data sets using *k*-means (Fig 8). The cluster maps (Fig 8A, B) exhibit clear signatures of inoculum effects, as once again the boundary between the “no growth” cluster and all other clusters is a function of both antibiotic concentration and inoculum, and not aligned with only MIC across all inocula.

**Fig 8.**
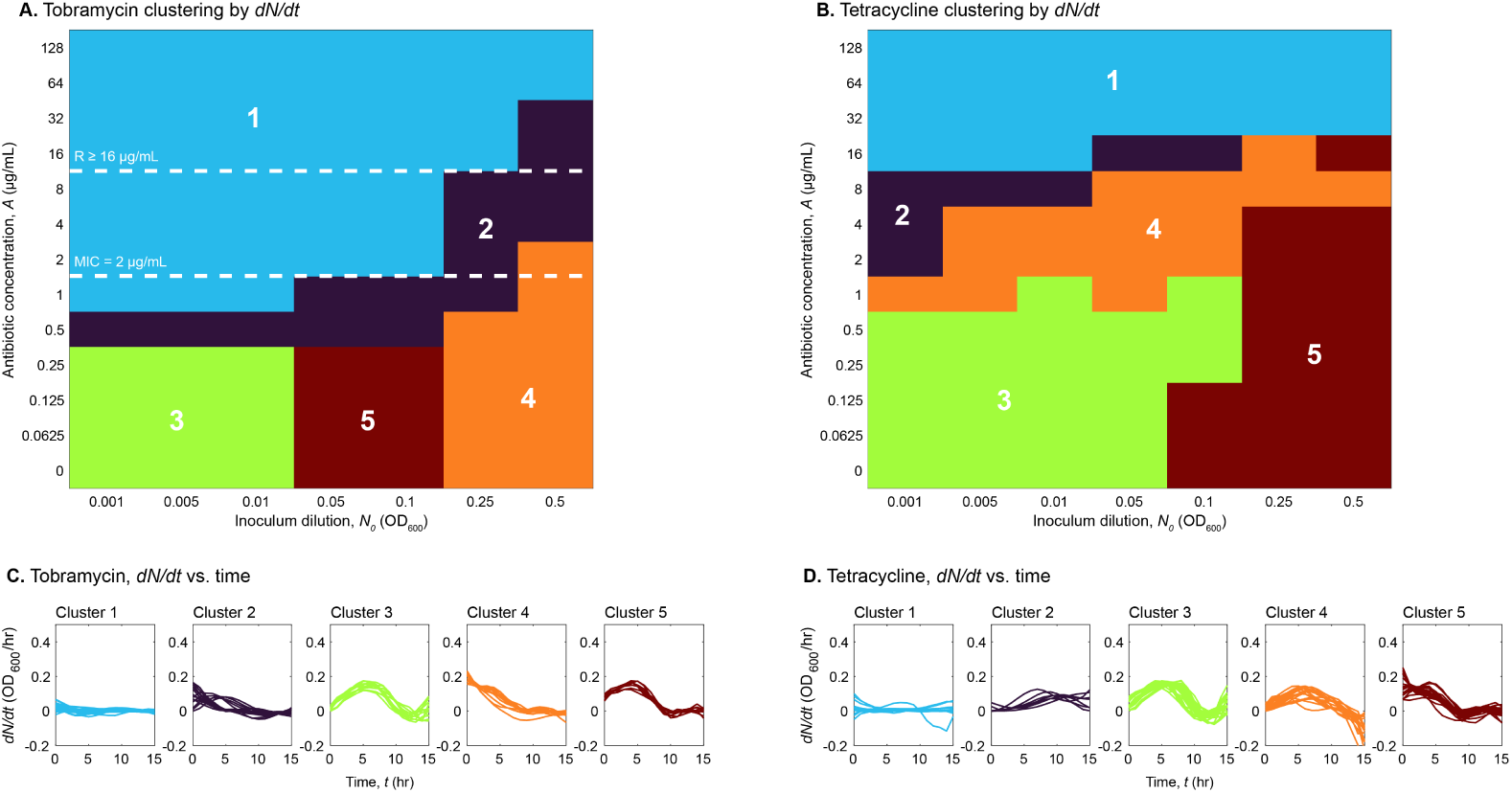
Results of clustering curves using *k*-means hold for bacterial growth dynamics when exposed to tobramycin and tetracycline. *k*-means clustering on the tobramycin and tetracycline data sets recovers five regimes determined jointly by antibiotic concentration *A* and inoculum *N*_0_. Colors and labels denote clusters in antibiotic dose, inoculum size treatment space. Panels A and B show the clusters of each treatment in antibiotic dose vs. inoculum space. The white dashed lines in Panel A denote MIC, labeled MIC = 2 µg*/*mL (Materials and methods) and the CLSI breakpoint for resistance of *P. aeruginosa* to tobramycin, labeled *R* ≥ 16 µg*/*mL [31]. These have not been included for tetracycline as it is not traditionally used clinically for treatment of *P. aeruginosa* infection. Panels C and D show corresponding approximate derivatives *dN/dt* vs. time for each cluster.

While some of our additional candidate inoculum effect models perform better overall when applied to the tobramycin and tetracycline data sets as compared to meropenem, the differences in average RMSE across the treatment space are small, suggesting that Eq 7 provides the best general model across drug, dose, and inoculum size. The relative success of other candidate models, however, does not refute any of our claims but rather continues to provide support for the presence of antibiotic-induced positive density dependence in *P. aeruginosa* population dynamics. Additionally, given differences in underlying mechanisms of action, it is reasonable that signatures of positive density dependent effects may vary across drugs, which can be observed by contrasting the weak Allee model parameters for meropenem, tobramycin, and tetracycline (Table 2).

## Discussion

Our findings provide quantitative insights into how inoculum size and antibiotic exposure combine to shape bacterial population dynamics. This work bridges ecology, quantitative modeling, and clinical microbiology by identifying patterns in bacteria-drug interactions that cannot be captured by standard antibiotic susceptibility metrics such as MIC. Below, we highlight the methodological, conceptual, and clinical relevance of our findings, and outline future directions.

### Quantitative modeling insights for microbial population dynamics

In this paper, we identify a composite model that recapitulates nonlinear and density dependent effects of antibiotic exposure and inoculum size by connecting mathematical models with experimental data.

Throughout our analysis, we consider different functional forms, algorithms for model fitting, and the number of parameters required for each model when attempting to find a differential equation-based model to describe combined antibiotic and inoculum effects. The goal was to identify models that were generalizable, biologically relevant, and grounded in experimental data, not just selected for their convenience or history of use. Given the wide variation in initial condition needed in our data set to investigate inoculum effects, we find that methods of model estimation relying on fitting a solution of an ODE to density data *N* (*t*) are insufficient to capture the diversity of dynamical behavior that we see in Fig 4.

Using Algorithm 1, a density-based fitting algorithm that doesn’t explicitly account for inoculum size, we observe a systematic underestimation of transient dynamics when antibiotic impacts are small and overestimation for treatments producing little to no growth. In this case, the optimization problem attempts to minimize error over time and for multiple growth regimes simultaneously, highlighting that even if inoculum size doesn’t impact dynamics explicitly, large variation in initial condition can skew model fitting and dominate over more interesting and important transient dynamics.

In contrast, Algorithm 2, which fits *dN/dt*, offers a substantial improvement. Here, we obtain a smaller parameter set, just the size of the number of model parameters, than when fitting each antibiotic case or each combined antibiotic dose, inoculum size treatment individually, as in Algorithms 1 or 3. Identifying a reduced set of parameters indicates that the model captures generalized mechanisms rather than responses specific to individual experimental settings. Furthermore, this approach can mitigate effects of wide variation in density by “aligning” our experimental trajectories according to their approximate derivative or net growth rate. Fig 4 provides a clear example of how considering derivative trajectories reduces variation in our data set as compared to density trajectories. While dynamics-based approaches have the added benefit of allowing us to capture multiple qualitative behaviors that vary based on initial conditions, they incur the added task of having to approximate the derivative. Even with hourly data, this can be noisy and lead to issues with fitting, emphasizing the need for increased temporal resolution when collecting biological data.

We conclude that fitting approximate derivatives to ODE models via “gradient matching” [37] type protocols is necessary for capturing diverse system dynamics, especially when densities vary over multiple orders of magnitude. We also emphasize that the type and resolution of data, choice of loss function to minimize, and the selection of bounds and initial guesses for parameters with biological relevance should all be considered and evaluated when utilizing model estimation methods.

### Ecological insights into pathogen clinical control and resistance metrics

Microbial populations experience stress as a result of environmental perturbations, invoking responses on cellular, population, and community levels. In the context of human infection, antibiotics are a common perturbation, either as the target of treatment or via bystander exposures [45–49]. Our results show how relying on standard, MIC-based summaries at a fixed inoculum can overestimate antibiotic impact on population dynamics. Our results can therefore help to explain instances of apparent mismatch between susceptibility tests and observed clinical treatment outcomes [50–52], even in the absence of more complex factors stemming from community ecological [47, 48] or evolutionary [49] processes.

In Fig 2, we clearly show that antibiotic exposure impacts both growth rate and yield, consistent with a recent study looking at how antibiotic exposure negatively impacts resource utilization [5]. Exposure to antibiotics presents a stressor to bacterial growth, leading to lower productivity [5]. Growth inefficiencies could result from bacterial strategies to prevent or minimize damage from stress or from the perturbing impact of metabolic imbalances on bacterial growth and yield [53, 54]. Additionally, while both time series data and ODE models assume homogeneity in antibiotic exposure and bacterial susceptibility, populations are exposed to heterogeneous antibiotic concentrations in space and time and are heterogeneously susceptible to antibiotic exposure [19, 55–59]. On a population scale, these effects are averaged: we see reduced growth yield as only a portion of the population is growing and survives. In Fig 2D-F, we show that antibiotic effects are nonlinear, flagging that linear approximations underestimate effects at low doses and overestimate effects at high doses.

We also investigated how bacterial density at the start of antibiotic treatment modulates antibiotic effects. Our results exhibit clear signatures of inoculum effects (Fig 3, Fig 4, and S4 Fig), where the functional MIC increases with inoculum size. We hypothesized that higher-order nonlinearities describing positive density dependence were responsible for this observed effect. Initial investigation using SINDy (Sparse Identification of Nonlinear Dynamics [60])—a symbolic regression algorithm that identifies polynomial terms and their coefficients—identified higher-order density terms consistent with this hypothesis, but lacked biological interpretation (just polynomial terms). By investigating the functional forms in our candidate models (Section 1.2 in S1 Text), we were able to propose and evaluate different potential mechanisms of density dependence, providing biological relevance and ultimately, identifying an antibiotic threshold dependent weak Allee effect as an adequate description of population dynamics (Fig 5, S5 Fig-S8 Fig).

Generally, the individual candidate models we tested struggled to capture the wide range of dynamics (no growth, linear trajectories to normal growth, S-shaped curves) without a “switch” where positive density dependent growth turns on. Our composite model (Eq 7) offers a possible strategy, as it combines the logistic growth model with saturating antibiotic loss with a weak Allee effect at higher antibiotic exposures. These conclusions are also generalizable to both tobramycin and tetracycline drug effects. By extending our investigation to these additional drugs, we highlight that this qualitative pattern of stress response in *P. aeruginosa* is more general than just in response to a certain drug or specific antibiotic mechanism of action.

A recurring theme in our work is the limitation of MIC as a susceptibility metric. To address the need for improved antibiotic susceptibility metrics, we point to several avenues raised by our work. Foremost, our mathematical model (Eq 7) introduces a novel parameter, *A_thresh_*, which defines the antibiotic dose threshold at which inoculum effects “switch on”. From a practical standpoint, *A_thresh_* helps to quantify the impact of inoculum effects on antibiotic susceptibility. In the case that *A_thresh_* is greater than the CLSI breakpoint, our model would predict no induction of positive density dependence with increasing inoculum size in the range of safe antibiotic dosage. Here, MIC may be a sufficient description of antibiotic susceptibility. However, in the case where *A_thresh_* is lower than the breakpoint, it indicates that inoculum effects need to be accounted for in determining susceptibility. We suggest that in this case, the clustering protocol employed in Fig 4, Fig 8, and S4 Fig may offer a dynamic-led strategy for determining antibiotic susceptibility, defined by the boundary between the “no growth” cluster (Cluster 1) and other clusters. While both our model and clustering approach require additional data, collection of OD data with sufficient time resolution is largely accessible with a simple plate reader. This provides a computationally simple strategy that leverages transient dynamics and inoculum size in determining antibiotic susceptibility.

### Limitations and future directions

Our approach prioritized experimental and mathematical model simplicity and tractability, and therefore presents a baseline for additional avenues of research. From an experimental standpoint, important future avenues include investigations on how population dynamics depend on bacterial species and strain ID, on growth media, and on opportunities for biofilm growth and other spatial structuring. We also flag that while the use of optical density measurements of bacterial densities allows for the generation of high-resolution time series data that is necessary for our approach, it is also vulnerable to systematic biases [61–63]. We note that alternate quantitation methods also face severe limitations, notably qPCR methods will count genomic material from cells killed by antibiotics [64], and CFU counting methods are too laborious to support high resolution time series estimation necessary for our gradient matching approach.

An additional future avenue is to pursue connections to the molecular mechanistic basis of weak Allee effects. A first step in that direction would be to transcriptomically profile populations under different antibiotic and inoculum conditions, to see both whether transcriptomic patterns reflect the population dynamical clusterings illustrated in Fig 4 and Fig 8, and whether specific profiles relate to established molecular mechanisms of drug susceptibility and inoculum effects.

## Conclusion

Data collection and resolution, *a priori* model forms and assumptions, and inference choices all impact our ability to efficiently describe the dynamics of microbial growth under variable antibiotic and density conditions. We conclude that bacterial growth under perturbation exhibits regimes of transient dynamical behavior, where most models can adequately capture low to no growth and near normal growth scenarios, but fail to capture intermediate dynamics unless an antibiotic threshold dependent weak Allee effect is introduced. By utilizing a clustering based approach on population dynamic trajectories, we are able to capture signatures of transient dynamics in these regimes—offering potential strategies to develop novel antibiotic susceptibility metrics that capture transient dynamical properties of pathogen-antibiotic interactions.

## Materials and methods

### Bacterial pre-culture and experimental treatment

We collect fine-scale temporal data of *P. aeruginosa* (PAO1) under a range of antibiotic exposures and with a range of inoculum concentrations. We collect bacterial density time curves for all combinations of *p* = 12 (meropenem) or *p* = 13 (tobramycin, tetracycline) antibiotic concentrations, *n* = 7 different inoculum sizes, and 3 antibiotic drugs (meropenem, tobramycin, and tetracycline).

*P. aeruginosa* strain PAO1 Nottingham was revived from frozen stock by streaking on LB agar plates and growing overnight at 37°C. A single colony was then cultured overnight in LB broth at 37°C with shaking. The culture was then washed in an equal volume of PBS and diluted in LB broth to an optical density of OD_600_ = 0.001 - 0.5 (referred to as the “inoculum dilution”). 10 µL were inoculated into appropriate wells of a 96-well plate containing 90 µL of various concentrations of antibiotic in LB broth.

Plates were incubated in a BioTek BioSpa 8 Automated Incubator (Agilent) and the OD_600_ was measured every hour in a BioTek Cytation 5 plate reader (Agilent).

### Antibiotic susceptibility assays and susceptibility determination

Wells of a 96-well plate were filled with either 100 µL blank media or 50 µL of tobramycin or meropenem diluted to appropriate concentrations in appropriate media. 50 µL aliquots of washed and diluted bacterial cultures were then added to antibiotic containing or blank wells. Plates were incubated at 37°C in a BioSpa 8 microplate automated incubator (Agilent) and OD_600_ was measured every 2 hours for 16 hours in a Cytation 5 plate reader (Agilent).

MIC for meropenem or tobramycin in LB (Fig 4 and 8) was defined as the lowest tested antibiotic concentration under which OD_600_ at 16 hours was ≤ 0.1 when grown using our lowest inoculum size, approximating standard protocols [25].

### Data availability and processing

Experimental data is available in supplemental files: S1 Data, S2 Data, and S3 Data. Bacterial density data was minimally processed for use in model fitting and classification (clustering). For all antibiotic and inoculum treatment combinations, time series were medium blank corrected, the first time point was omitted as an artifact of data collection, and negative OD_600_ values were set to 0.0001. Time series were averaged across replicates.

The code for analysis was created using MATLAB R2021b–academic use and R2024a. Figures were produced using MATLAB R2021b–academic use and R2024a, and Adobe Illustrator 2023.

### Model fitting

Using the processed data (only the first 15 hours to focus on growth dynamics), we parameterized the models defined in the main text and Section 2 of S1 Text using regression following from commonly used approaches [36–38, 65] over a variety of assumptions, constraints, and degrees of freedom. We utilize three algorithms for fitting data:

- *Algorithm 1.* Models are fit for each antibiotic condition separately, using density *N* (*t*), and assuming variation in inoculum size has no effect on dynamics. This fitting protocol results in a parameter set for each of the *p* antibiotic concentrations tested in each data set.
- *Algorithm 2.* Models are fit for all experimental treatments simultaneously using approximate derivatives, *dN/dt*. This fitting protocol results in a single parameter set for each data set.
- *Algorithm 3.* Models are fit for all experimental treatments (antibiotic dose and inoculum size combination) separately, using density *N* (*t*). This fitting protocol results in a parameter set for each of the *T* treatments (*T* = *p* antibiotic concentrations times *n* inoculum sizes).

Algorithms 1 and 2 are presented in our results as a part of our final analyses and computational pipeline. Algorithm 3 was used to initially understand the dynamics of our data set, in development of our final computational pipeline, and to select the values of growth rate *r* and carrying capacity *k* in the absence of antibiotic exposure for use with Algorithms 1 and 2. We further describe the data fitting protocols and models used in S1 Text. The code is publicly available at: https://github.com/GaTechBrownLab/abx-inoculum-effects.

### Clustering

Starting with the data from *P. aeruginosa* exposed to meropenem, we cluster the approximate derivative *dN/dt* using *k*-means [43]. We use kmeans in MATLAB [66] and select the number of clusters using evalclusters by evaluating clustering using up to 6 clusters (rule of thumb, 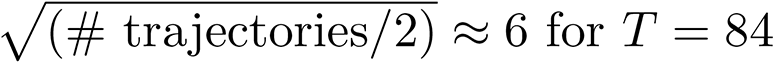 treatments in S1 Data), and selecting the optimal number based on the silhouette criterion. The silhouette criterion measures the similarity of a data point to points in the same cluster as compared to points in other clusters [67]. In the case of meropenem, the optimal number of clusters is 5. We cluster all antibiotic data sets independently, but apply the number of clusters defined for meropenem to the tobramycin and tetracycline data sets. The cluster colors and numbering are applied such that Clusters 1-5 in Fig 4 and Fig 8 capture similar dynamics across the independent clusterings. We follow the same protocol with bacterial density *N* (*t*), where we find the optimal number of clusters for meropenem is 2 (S4 Fig).

## Acknowledgments

We thank the Brown Lab and the Center for Microbial Dynamics and Infection at Georgia Tech for valuable discussion and comments on earlier drafts.

We thank the following organizations for funding: the National Science Foundation, award NSF 2406985 (to S.P.B. and R.K.); the Cystic Fibrosis Foundation, award BROWN23I0 (to S.P.B.); the Department of Defense, through the National Defense Science and Engineering Graduate Fellowship Program (S.S.); and the Achievement Rewards for College Scientists Foundation Atlanta Chapter (S.S.). Edits were completed while the author (S.S.) held an NRC Research Associateship award at the United States Naval Research Laboratory.

## Supporting information

**S1 Fig. Experimental time series data by meropenem concentration.** Averaged trajectories from experimental treatment of *P. aeruginosa* with varied meropenem dose and inoculum concentration. Colors denote inoculum dilution.

**S2 Fig. Effective growth rate and growth yield do not vary with *N*_0_ in Eq 5.** Simulation of Eq 5 with *A* = 0.5 µg*/*mL (Panels A-C) and *A* = 32 µg*/*mL (Panels D-F). Panels A and D: bacterial population density tracked over time starting from varied initial density (low inoculum = dark blue, high inoculum = dark red). Panels B and E: corresponding effective growth rate *r*(*A*) vs. initial density *N*_0_. Panels C and F: corresponding growth yield *N* ^∗^ vs. initial density *N*_0_. Parameters: *r* = 1.4052, *k* = 1.1249, *l* = 1.1764, *h* = 1.2117, *A*_50_ = 1.0537, *A* = 0.5, 32, *N*_0_ = [0.001, 0.005, 0.01, 0.05, 0.1, 0.25, 0.5], Δ*t* = 0.01.

**S3 Fig. Existence of inoculum effects in experimental data under low and high antibiotic concentrations.** Panels A-D: experimental data for bacterial density time series with no antibiotic (*A* = 0 µg*/*mL, black lines) and low antibiotic exposure (*A* = 0.25 µg*/*mL, red lines). Panels E-H: experimental data for bacterial density time series with no antibiotic (*A* = 0 µg*/*mL, black lines) and high antibiotic exposure (*A* = 16 µg*/*mL, red lines). Inoculum size increases left to right identically from Panel A to D and E to H, where *N*_0_ = 0.001, 0.01, 0.1, 0.5.

**S4 Fig. Density curves clustered using *k*-means show two regimes of dynamic behavior as decided by combinations of both *A* and *N*_0_.** Colors and labels denote clusters in (antibiotic, inoculum)-treatment space. White dashed lines denote MIC determined via standard antibiotic susceptibility testing, labeled MIC = 2 µg*/*mL (Materials and methods) and the CLSI breakpoint for resistance of *P. aeruginosa* to meropenem, labeled *R* ≥ 8 µg*/*mL [31]. Panels B and C show corresponding population density time series for each cluster.

**S5 Fig. Model vs. data by meropenem concentration for fitting the logistic growth model with saturating antibiotic loss** (Eq 5) using Algorithm 2. Colors denote inoculum dilution where dark blue is low inoculum, dark red is high inoculum. Parameters are defined in Fig 2. Corresponding RMSE shown in Fig 5A and Panel B of S7 Fig.

**S6 Fig. Model vs. data by meropenem concentration for fitting the antibiotic threshold dependent weak Allee effect model (Eq 7) using Algorithm 2**. Colors denote inoculum dilution where dark blue is low inoculum, dark red is high inoculum. Parameters are defined in Table 2 for meropenem. Corresponding RMSE shown in Fig 5B.

**S7 Fig. Heat maps of RMSE for data from all treatments (antibiotic concentration and inoculum size combination) for each candidate model.** RMSE for each candidate model fit using Algorithm 2 (see Section 2.3 in S1 Text for additional model details). Panel A: logistic growth model with linear antibiotic loss (Eq 2 with *L*(*A*) = *L_lin_*(*A*), Eq 3). Panel B: logistic growth model with saturating antibiotic loss (Eq 5). Panel C: Allee effect model (Eq S.3). Panel D: cooperation model (Eq S.4). Panel E: effective antibiotic model (Eq S.5). Panel F: expanded logistic growth model (Eq S.6). Parameters for Panels A and B follow from Fig 2. Parameters for Panels C-F are defined in Section 1.2 of S1 Text.

**S8 Fig. Model vs. data by meropenem concentration for fitting each candidate model using Algorithm 2.** Colors denote inoculum dilution where dark blue is low inoculum, dark red is high inoculum. Antibiotic concentrations: 0.125, 2, 32 µg*/*mL (time vs. density axes by row, left to right). Panel A: logistic growth model with linear antibiotic loss (Eq 2 with *L*(*A*) = *L_lin_*(*A*), Eq 3). Panel B: Allee effect model (Eq S.3). Panel C: cooperation model (Eq S.4). Panel D: effective antibiotic model (Eq S.5). Panel E: expanded logistic growth model (Eq S.6. Parameters for Panel A are follow from Fig 2. Parameters for Panels B-E and additional model details are defined in Section 1.2 of S1 Text. Corresponding RMSE shown in S7 Fig.

**S1 Text. Mathematical modeling and computational pipelines.**

**S1 Data. Experimental data for *P. aeruginosa* exposed to meropenem.**

**S2 Data. Experimental data for *P. aeruginosa* exposed to tobramycin. S3 Data. Experimental data for *P. aeruginosa* exposed to tetracycline.**

## Notes

https://github.com/GaTechBrownLab/abx-inoculum-effects

### Competing Interest Statement

The authors have declared no competing interest.

### Summary of Updates

Text S1 updated; Github repository information included.

