## Supplementary material for "Quantifying antibiotic susceptibility and inoculum effects using transient dynamics of *Pseudomonas aeruginosa*": S1 Text

### S1 Text. Mathematical modeling and computational pipelines.

### 1 Mathematical models

#### 1.1 Baseline models: logistic growth model with antibiotic effects

We define the classic logistic growth model in Eq 1. The model has two equilibrium points, $N^* = 0$ and $N^* = k$, where the population either tends to extinction (if $r < 0$) or the carrying capacity (if $r > 0$), respectively, as $t \to \infty$. In the event that $r = 0$, the population will remain at its initial density, $N_0$. Here, the maximal growth yield is akin to the carrying capacity and subsequently, the equilibrium point $N_2^*$, where the population survives.

To model the effect of antibiotics, we use $L(A)$ as an initial approximation to describe the rate of population loss due to antibiotic exposure $A$. We consider two approaches: (I) antibiotics modify the maximal growth rate, leading us to replace the maximal growth rate in the absence of antibiotic exposure, $r$, in Eq 1 with a function of antibiotic concentration, which we refer to as the effective growth rate, $r(A) = r - L(A)$; and (II) antibiotics modify both the growth rate and yield, which we capture through an additional antibiotic concentration dependent loss term, $L(A)N$ (Eq 2).

In approach I, we define

$$\frac{dN}{dt} = r(A)N\left(1 - \frac{N}{k}\right) = r(A)N - \frac{r(A)}{k}N^2 \tag{S.1}$$

and again find two equilibrium points: $N^*_{I,1} = 0$ and $N^*_{I,2} = k$, where now the population tends to extinction if $r(A) < 0$ or carrying capacity if $r(A) > 0$, respectively. The maximal growth yield continues to be the carrying capacity and is not modified by antibiotic exposure.

In approach II, we define

$$\frac{dN}{dt} = rN\left(1 - \frac{N}{k}\right) - L(A)N = \big(r - L(A)\big)N - \frac{r}{k}N^2 = r(A)N - \frac{r}{k}N^2$$

(Eq 2 in the main text), where now the equilibrium points are $N^*_{II,1} = 0$ and $N^*_{II,2} = \frac{k}{r}(r - L(A))$. Similarly to Eq S.1, the population tends to extinction if $r(A) < 0$ and maximal growth yield $N^*_{II,2}$ if $r(A) > 0$. We note that the linear coefficient in Eq 2 is consistent with the linear coefficient (i.e. the effective growth rate $r(A)$) in Eq S.1 above, but now the maximal growth yield is dependent on the antibiotic dose such that the population cannot reach carrying capacity $k$ given antibiotic exposure. Given that our experimental data exhibits population declines in both growth rate and yield as a function of antibiotic exposure (Fig 2D, S1 Fig), we select the approach II model (Eq 2) as our starting point. In the main text, we explore two functional forms of the antibiotic loss rate, $L(A) = L_{lin}(A)$ (Eq 3) and $L(A) = L_{sat}(A)$ (Eq 4), leading to the selection of our baseline model, the logistic growth model with saturating loss,

$$\frac{dN}{dt} = rN\left(1 - \frac{N}{k}\right) - l\frac{A^h}{A_{50}^h + A^h}N$$

(Eq 5 in the main text). We define the net growth rate for Eq 5,

$$g(N) = r\left(1 - \frac{N}{k}\right) - l\frac{A^h}{A_{50}^h + A^h}, \tag{S.2}$$

and note that net growth is linear with respect to density $N$ and nonlinear with respect to antibiotic dose $A$. Given signatures of inoculum effects, antibiotic-induced positive density dependence, in our experimental data set (Fig 3, S3 Fig), we propose a series of candidate models which introduce nonlinear density dependence not captured in Eq 5 and S.2.

#### 1.2 Candidate models

We analyzed additional mathematical models and fit them to our experimental data using the data fitting protocols and evaluation methods described in Sections 2-4. Models were selected based on their functional forms and relevance for describing potential biological mechanisms of inoculum effects. Section 1.2.5 describes our proposed composite model, Eq 7.

##### 1.2.1 Allee effect model

We define the Allee effect model [1, 2],

$$\frac{dN}{dt} = \left[rN\left(1 - \frac{N}{k}\right) - l\frac{A^h}{A_{50}^h + A^h}N\right]\left(\frac{N-a}{k}\right) \quad \text{(S.3)}$$

where we introduce a density-threshold dependent Allee effect term, $(N - a)/k$, with $a$ denoting the density threshold. The model captures negative density dependent growth when population size is small ($N < a$) and positive density dependent growth when the population size is large ($N > a$). In addition to the equilibrium points for Eq 5 ($N_1^* = 0$ and $N_2^* = \frac{k}{r}(r - L_{sat}(A))$), the Allee effect model has an additional equilibrium point at the density threshold, $N_3^* = a$. Net growth is defined, $g(N) = [r(1 - \frac{N}{k}) - L_{sat}(A)](\frac{N-a}{k})$, showing that net growth is a nonlinear function of both density and antibiotic concentration.

The modifications of Eq 5 in Eq S.3 lead to an additional equilibrium point, allowing for potential bistability; and increased polynomial order of growth, carrying capacity limitation, and antibiotic loss terms, describing higher order nonlinearities in these processes. We hypothesized that Eq S.3 would capture signatures of inoculum effects in the form of growth bistability, where growth yield depends on initial density, dependent on a threshold [3–5].

Panel C in S7 Fig and Panel B in S8 Fig show the RMSE and sample time series trajectories for Eq S.3. Parameters from fitting via Algorithm 2 are: $r = 1.4052$, $k = 1.1249$, $l = 0.8218$, $h = 1.0832$, $A_{50} = 1.1238$, $a = 0.1162$.

##### 1.2.2 Cooperation model

We define the cooperation model, adapted from the model concept in [6],

$$\frac{dN}{dt} = rN\left(1 - \frac{N}{k}\right) - l\frac{A^h}{A_{50}^h + A^h}N + bN^2 \quad \text{(S.4)}$$

where $b$ describes the cooperative benefit from intraspecific interactions. We modify Eq 5 including a quadratic growth benefit term representing cooperativity, that specifically modifies the net intraspecific interaction rate. The equilibrium points are: $N_1^* = 0$ and $N_2^* = (r - L_{sat}(A))/(\frac{r}{k} - b)$. We can define net growth as, $g(N) = r(1 - \frac{N}{k}) + bN - L_{sat}(A)$. Net growth is a linear function of density $N$ and a nonlinear function of antibiotic concentration $A$.

Cooperative behaviors offering population level protection from antibiotics via mechanisms such as: spatial structuring, degradation, public good production, and quorum sensing [7, 8] provide a possible mechanism for inoculum effects, which we capture generally in Eq S.4.

Panel D in S7 Fig and Panel C in S8 Fig show the RMSE and sample time series trajectories for Eq S.4. Parameters from fitting via Algorithm 2 are: $r = 1.4052$, $k = 1.1249$, $l = 0.9703$, $h = 0.6000$, $A_{50} = 0.5797$, $b = 0.1702$.

##### 1.2.3 Effective antibiotic model

We define the effective antibiotic model, adapted from [9],

$$\frac{dN}{dt} = rN\left(1 - \frac{N}{k}\right) - l\frac{(\frac{A}{N})^h}{A_{50}^h + (\frac{A}{N})^h}N \quad \text{(S.5)}$$

where we substitute antibiotic concentration $A$ in Eq 5 with $A/N$. Antibiotic concentration is now linearly scaled by density. We again have an extinction equilibrium point $N_1^* = 0$, and additional equilibria $N^*$ that are solutions to $r(1 - \frac{N^*}{k}) - L_{sat}(\frac{A}{N^*}) = 0$. Net growth is defined, $g(N) = r(1 - \frac{N}{k}) - L_{sat}(\frac{A}{N})$ and is a nonlinear function of density $N$ and antibiotic concentration.

Biologically, Eq S.5 captures a titration effect, or a lower number of binding sites per binding targets, which is a popular theory for the mechanism underlying inoculum effects: more bacterial cells means less antibiotic per cell [10, 11].

Panel E in S7 Fig and Panel D in S8 Fig show the RMSE and sample time series trajectories for Eq S.5. Parameters from fitting via Algorithm 2 are: $r = 1.4052$, $k = 1.1249$, $l = 0.9527$, $h = 0.7393$, $A_{50} = 1.8948$.

###### 1.2.4 Expanded logistic model

The expanded logistic model is defined,

$$\frac{dN}{dt} = rN^u\left(1 - \frac{N}{k}\right)^v - l\frac{A^h}{A_{50}^h + A^h}N \tag{S.6}$$

where we introduce two additional parameters describing growth and resource utilization scaling: $u$ is the growth regulator, and $v$ is the depletion factor [12, 13]. As $u$ and $v$ define the polynomial order of our model, we will have $u + v$ equilibrium points defined as the values $N^*$ that satisfy $dN/dt = 0$. We define net growth as, $g(N) = rN^{u-1}(1 - \frac{N}{k})^v - L_{sat}(A)$, showing nonlinear density and antibiotic dependence.

Eq S.6 leads to modification of growth rate and yield, and $u$ and $v$ change the polynomial order of growth and carrying capacity limitation terms, introducing nonlinear dynamics. The expanded logistic model is used to approximate a growth delay or lag [13]—in line with idea that resistant sub-populations or persistence may be driving inoculum effects [14–17].

Panel F in S7 Fig and Panel E in S8 Fig show the RMSE and sample time series trajectories for Eq S.6. Parameters from fitting via Algorithm 2 are: $r = 1.4052$, $k = 1.1249$, $l = 0.8484$, $h = 1.1210$, $A_{50} = 0.7970$, $u = 1.1477$, $v = 1$.

###### 1.2.5 Proposed composite model: antibiotic threshold dependent weak Allee effect model

Based on the success of Eq S.3 at high inoculum sizes and high antibiotic concentrations, we explored additional functional forms of Allee effects [1, 18, 19], their impacts to growth and antibiotic loss components of the model, and antibiotic threshold-dependent behavior. Combined with the success of Eq 5 at low inoculum sizes and low antibiotic concentrations, we found that a composite model—integrating Eq 5 at low antibiotic concentrations with a weak Allee effect at high antibiotic concentrations—was able to preserve features of each model in different dynamical regimes. To define the composite model, we introduced a "switch", where a weak Allee effect modifying only the growth terms (and not the antibiotic loss rate) is present above a critical antibiotic threshold, $A_{thresh}$. While our initial investigation characterized a strong Allee effect (Eq S.3), we found that a weak Allee effect better approximated our experimental observations. In contrast to strong Allee effects, which lead to negative per-capita net growth below a threshold, a weak Allee effect encompasses positive per-capita net growth deviating from logistic growth, where the magnitude (and not the sign) of the effect is dependent on the relation of the population size to the Allee threshold $a$ [18, 19].

We define our composite model in Eq 7 in the main text. When $A \leq A_{thresh}$, we recover the dynamics and long-time behavior of Eq 5. When $A > A_{thresh}$, net growth is nonlinear with respect to density, defined $g(N) = r(1 - \frac{N}{k})f(N) - L_{sat}(A)$, as a result of the weak Allee effect.

#### 2 Model fitting protocols

In order to explore the impacts of antibiotic concentration and inoculum size on population dynamics and model parameters, we defined three algorithms for fitting models to our experimental data. By comparing and contrasting the different data fitting protocols described below, we not only aimed to identify a general model of bacterial population growth and inoculum effects given antibiotic exposure from our data set, but also investigated best practices for model fitting to data sets with large variation in initial condition.

#### 2.1 Notation

We define bacterial density given time $\mathbf{t} = [t_0, t_1, ..., t_m]^\intercal$ and antibiotic concentration $A$ as an $m \times n$ matrix, $\mathbf{N}_A$, where $m$ is the number of time points and $n$ is the number of inoculum conditions. We measure time in hours (hr), antibiotic concentration in µg/mL, and density as $OD_{600}$. We test $p = 12$ antibiotic concentrations, $A \in [0, 128]$ µg/mL (for meropenem; $p = 13$ in the same range for tobramycin and tetracycline), and $n = 7$ inoculum conditions, corresponding to inoculum dilutions: $i_k = [0.001, 0.005, 0.01, 0.05, 0.1, 0.25, 0.5]$ for $k = 1, ..., n$ (where $i_k \approx \mathbf{N}_A(0, i_k)$, referred to as $N_0$ in the context of our models described above and in the main text). For example, $\mathbf{N}_{A=0.125}(\mathbf{t}, i_1)$ is the time series corresponding to the antibiotic-inoculum treatment: $A = 0.125$ µg/mL of meropenem with inoculum dilution 0.001. We define the total number of treatments, where a treatment is the combination of one antibiotic dose and one inoculum condition, as $T = p \cdot n$.

#### 2.2 Algorithm 1

In Algorithm 1, we fit a given model using bacterial density $N(t)$ for each antibiotic treatment, yielding parameters that are a function of antibiotic concentration $A$ (e.g., for Eq. 1, $r(A)$ and $k(A)$). This results in a large parameter set of size equal to the number of model parameters times $p$ antibiotic concentrations to describe the whole data set. While here we assume that there is no explicit inoculum effect, we use the data from all $n$ inoculum size treatments, which can be interpreted as natural variation in the initial population size. The assumptions made by Algorithm 1 approximate traditional approaches to quantifying antibiotic susceptibility, such as MIC, that do not account for inoculum effects.

**Algorithm 1** Fitting the density by antibiotic concentration

For a given antibiotic concentration $A$, we fit the solution of a chosen model equation ($\hat{\mathbf{N}}_A$) to experimental data ($\mathbf{N}_A$) via a constrained least-squares optimization problem (using `lsqnonlin` in MATLAB [20] with objective function $\mathrm{SSE}_A$) to obtain effective parameters that are a function of $A$.

**for** $A$ **do**

**Step 1.** Define $\mathbf{N}_A = \begin{bmatrix} N_A(t_0, i_1) & N_A(t_0, i_2) & \dots & N_A(t_0, i_n) \\ N_A(t_1, i_1) & N_A(t_1, i_2) & \dots & N_A(t_1, i_n) \\ \vdots & \vdots & \ddots & \vdots \\ N_A(t_m, i_1) & N_A(t_m, i_2) & \dots & N_A(t_m, i_n) \end{bmatrix}$

**Step 2.** Pick initial parameter guess

**Step 3.** Solve chosen model ODE for $\hat{\mathbf{N}}_A$ using current parameters

**Step 4.** Calculate the sum of squares error over $m$ time points for the $k$th inoculum,

$$\mathrm{SSE}_k = \sum_{j=0}^{m} \left[ \hat{\mathbf{N}}_A(t_j, i_k) - \mathbf{N}_A(t_j, i_k) \right]^2$$

then sum across the inoculum condition: $\mathrm{SSE}_A = \sum_{k=1}^{n} \mathrm{SSE}_k$; we look to minimize $\mathrm{SSE}_A$

**Step 5.** Update parameter values using the Levenberg-Marquardt algorithm [21, 22]

**Step 6.** Iterate through Steps 3-5 until $\mathrm{SSE}_A$ is minimized

**Step 7.** Save the current parameter values

**end for**

#### 2.3 Algorithm 2

In Algorithm 2, we fit a given model using the derivative of bacterial density $dN/dt$, where antibiotic concentration $A$ is an independent variable, for the whole data set simultaneously. In contrast to previous work where antibiotic susceptibility parameters like MIC and $A_{50}$ are functions of inoculum size [23], we tackle variation in initial condition by fitting model dynamics versus model densities. This "gradient matching" approach [24], avoids biases in attempting to minimize error between the model prediction and experimental data for both low and high initial densities simultaneously, instead focusing on density dependencies in the dynamics. Algorithm 2 has the added benefit of producing a single parameter set of size equal to the number of model parameters.

For Algorithm 2, we approximate the derivative of our experimental data for a given treatment $N_A(\mathbf{t}, i_k)$ using the `gradient` function in MATLAB [20], verifying agreement with spline fitting from [25]. We define this approximation as $F_A(\mathbf{t}, i_k)$, and subsequently can define approximate net growth $g_A(\mathbf{t}, i_k) = F_A(\mathbf{t}, i_k)/N_A(\mathbf{t}, i_k)$. We stack the approximate derivatives from all $T$ treatments ($T = p$ antibiotic concentrations times $n$ inoculum sizes) in a $(m \cdot T) \times 1$ vector $\mathbf{F}$ (or $\mathbf{g}$ for net growth). We similarly define $(m \cdot T) \times 1$ vectors for time $\mathbf{t_{all}} = [\mathbf{t}_1, \mathbf{t}_2, ..., \mathbf{t}_T]^\intercal$ and antibiotic concentration $\mathbf{A_{all}} = [\mathbf{A}_1(\mathbf{t}), \mathbf{A}_2(\mathbf{t}), ..., \mathbf{A}_T(\mathbf{t})]^\intercal$, where $\mathbf{A}_a(\mathbf{t})$ is the antibiotic concentration of treatment $a = 1, 2, ..., T$ at time $t$ defined as constant in time or by experimental or simulated data. Then, we fit either the derivative $F(N) = dN/dt$ or net growth rate ($g(N) = \frac{1}{N}\frac{dN}{dt} = \frac{F(N)}{N}$, Eq 8). We define the predicted derivative and net growth values as $\hat{\mathbf{F}}$ and $\hat{\mathbf{g}}$, respectively. We utilize mean absolute error (MAE) ($l_1$ loss), as it generally prevents us from getting stuck at bounds and initial conditions as a result of local minima in the case of our candidate models and data.

**Algorithm 2** Fitting the dynamics of all antibiotic-inoculum treatments simultaneously

For a given data set, we fit a chosen model equation $F(N) = dN/dt$ (predicted $\hat{\mathbf{F}}$) to the derivative approximation from experimental data ($\mathbf{F}$) via a constrained optimization problem (using `fmincon` in MATLAB [20] with objective function $\text{MAE}_{tot}$) to obtain a single parameter set with independent variables $A$ and $t$. We outline the protocol for the derivative $F(N)$, but note that we can use the same protocol with net growth instead fitting Eq 8 (predicted $\hat{\mathbf{g}}$) to the approximation from experimental data ($\mathbf{g}$).

**Step 1.** Define $\mathbf{t_{all}}$ and $\mathbf{A_{all}}$, and approximate $\mathbf{F}$ from experimental data, as discussed above

**Step 2.** Pick initial parameter guess

**Step 3.** Calculate $\hat{\mathbf{F}}$ using current parameters

**Step 4.** Calculate the mean absolute error summed over $M = m \cdot T$ data points ($m$ time points $\times$ $T$ treatments),

$$\text{MAE}_{tot} = \frac{1}{M}\sum_{j=1}^{M}\left|\hat{\mathbf{F}}_{j,1} - \mathbf{F}_{j,1}\right|$$

We look to minimize $\text{MAE}_{tot}$

**Step 5.** Update parameter values using a Sequential Quadratic Programming (SQP) method [26]

**Step 6.** Iterate through Steps 3-5 until $\text{MAE}_{tot}$ is minimized

**Step 7.** Save the current parameter values

#### 2.4 Algorithm 3

In our final fitting protocol, Algorithm 3, we fit a given model using bacterial density $N(t)$ for each combined antibiotic dose $A$ and inoculum size $N_0$ treatment. Similar to Algorithm 1, we obtain a large parameter set of size equal to the number of model parameters times $T$ treatments ($T = p$ antibiotic concentrations times $n$ inoculum sizes) where fit parameters are now a function of both $A$ and $N_0$. This approach allows us to look at the effect of both antibiotic concentration and inoculum size on parameters and provides better model accuracy; however, it fails to provide a general functional form of the dynamics in the whole data set without additional data, model fitting, or model identification to define antibiotic dose and inoculum size dependencies.

**Algorithm 3** Fitting the density by antibiotic-inoculum treatment

For a given antibiotic concentration $A$ and inoculum dilution $i_k$, we fit the solution of a chosen model equation ($\hat{\mathbf{N}}_A(\mathbf{t}, i_k)$) to experimental data ($\mathbf{N}_A(\mathbf{t}, i_k)$) via a constrained least-squares optimization problem (using `lsqnonlin` in MATLAB [20] with objective function $\text{SSE}_k$) to obtain effective parameters that are a function of $A$ and $i_k$.

**for** $A$ **do**

**for** the $k$th inoculum dilution **do**

**Step 1.** Define $\mathbf{N}_A(\mathbf{t}, i_k) = \begin{bmatrix} N_A(t_0, i_k) \\ N_A(t_1, i_k) \\ \vdots \\ N_A(t_m, i_k) \end{bmatrix}$

**Step 2.** Pick initial parameter guess

**Step 3.** Solve chosen model ODE for $\hat{\mathbf{N}}_A(\mathbf{t}, i_k)$ using current parameters

**Step 4.** Calculate the sum of squares error $\text{SSE}_k$ over $m$ time points as defined in Algorithm 1; we look to minimize $\text{SSE}_k$

**Step 5.** Update parameter values via the Levenberg-Marquardt algorithm [21, 22]

**Step 6.** Iterate through Steps 3-5 until $\text{SSE}_k$ is minimized

**Step 7.** Save the current parameter values

**end for**

**end for**

#### 2.5 Success of derivative-based model fitting and full computational pipeline

Overall, we found that Algorithm 2 provides the best approach for efficiently capturing the combined impacts of antibiotic concentration and inoculum size on bacterial population growth. Given the diversity of population growth and no growth dynamics observed in our data sets (S1 Fig, Fig 4, Fig 8), selecting parameters based on $N(t)$ leads to a model that "averages out" these behaviors—systematically underestimating near normal growth dynamics (Clusters 3-5 in in Fig 4) and overestimating no to low growth dynamics (Cluster 1 in Fig 4)—as the algorithm tries to minimize error across all treatments and time. By using the rate of change in density $dN/dt$, we avoid biases in minimizing error over time, antibiotic concentration, and inoculum size.

Nevertheless, each algorithm has it strengths. By utilizing all three protocols in tandem, we were able to gain a full understanding of the dynamics and parameter dependencies in our experimental data sets, ultimately leading to the selection of a minimal model that produced results consistent with more complex forms (e.g., larger parameter set).

### 3 Parameter selection

For the parameterization of candidate models using Algorithm 2, we fixed the maximal growth rate $r$ and carrying capacity $k$ based on fitting Eq 1 for the no antibiotic case ($A = 0$) using Algorithm 3 and taking the average over the number of inoculum conditions. Table 2 contains these values of $r$ and $k$ for each data set. In the case of tobramycin and tetracycline, we use the same data for fitting the no antibiotic case as experiments were completed at the same time using the same bacterial stock. Our qualitative results were conserved when other protocols for fitting $r$ and $k$ were applied, as well as when $r$ and $k$ were left as open parameters in fitting the candidate models using Algorithms 1-3. Further, we confirmed that the selected values of $r$ corresponded to the range of doubling times observed experimental for *P. aeruginosa* [27]. All other parameters were left "open" for selection via model fitting within appropriate bounds.

Initial guesses and upper and lower bounds for the parameters used in fitting the candidate models were chosen to reflect biologically relevant parameters from the literature (MIC range [28,29] and growth rate range [27]), and estimates from our experimental data and initial fitting (difference in $OD_{600}$ between initial and final time points, final density, estimation of $h$ and $A_{50}$ using fit parameters for $r(A)$ and $k(A)$ in Fig 2E-F (green curves)). Baeder and Regoes (2019) provide a mathematical definition for relating pharmacodynamic parameters (MIC, $A_{50}$, and $h$) to compare our antibiotic loss parameters with experimentally measured values [23].

### 4 Model Evaluation

In order to compare model parameterizations, we utilize root mean squared error (RMSE) in order to quantify model accuracy in describing dynamics of the bacterial population density $N(t)$. RMSE describes the distance between our model predictions and observed data for a given antibiotic-inoculum treatment, and is defined

$$\text{RMSE} = \sqrt{\frac{\sum_{j=1}^{m}(N(t_j) - \hat{N}(t_j))^2}{m}}$$

where $N(t)$ are the observed (true) values, $\hat{N}(t)$ are the predicted values, and $m$ is the total number of time points (Eq 6 in the main text). We look for low RMSE. Given that we parameterize an ODE model, we use numerical simulation of $d\hat{N}/dt$ using the forward Euler method to solve for the solution $\hat{N}(t)$ of a given model and treatment. Fig 5 and S7 Fig depict heat maps of RMSE across the (antibiotic, inoculum)-space.
