## Supplementary figures and images for "Quantifying antibiotic susceptibility and inoculum effects using transient dynamics of *Pseudomonas aeruginosa*"

### S1 Fig

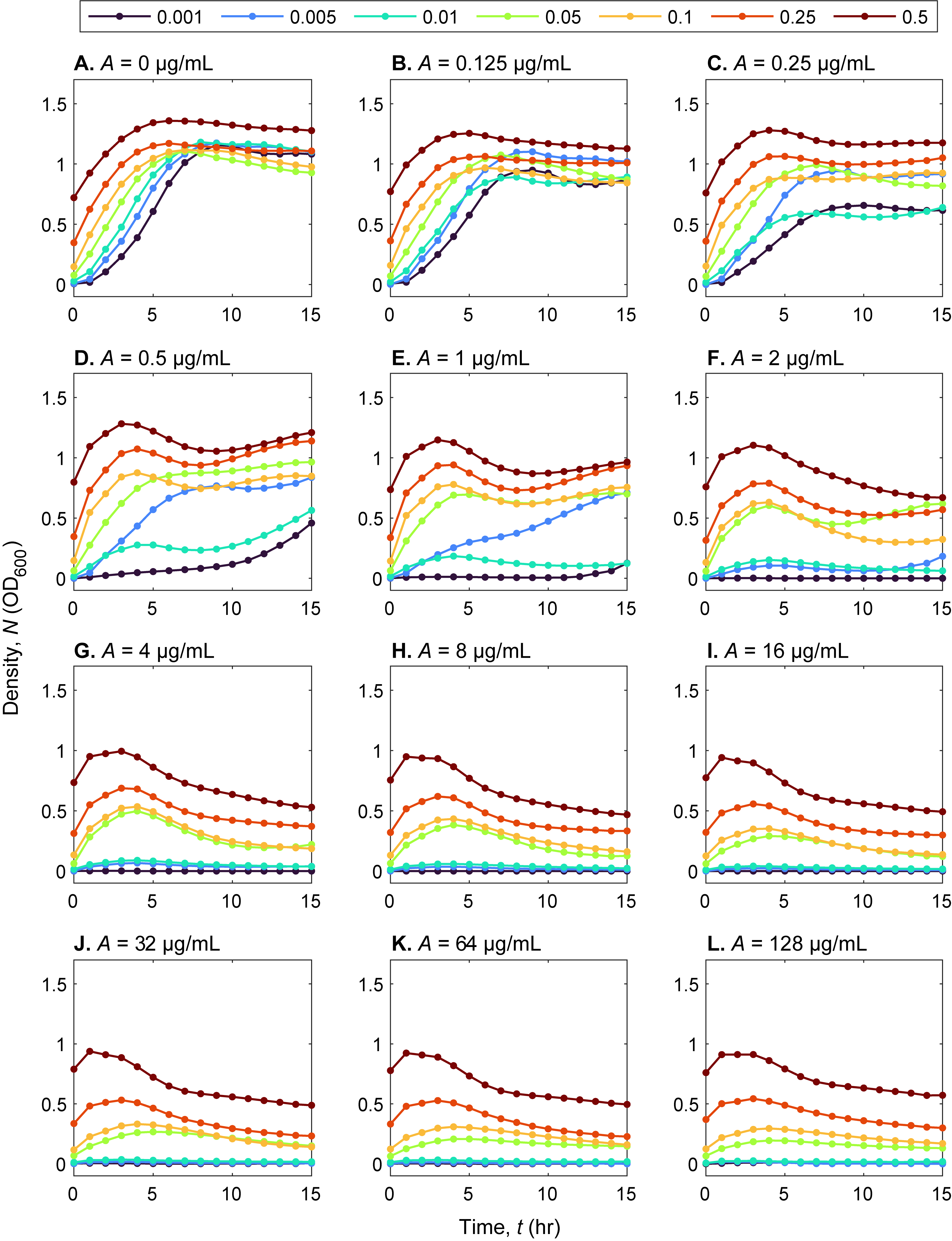

### S2 Fig

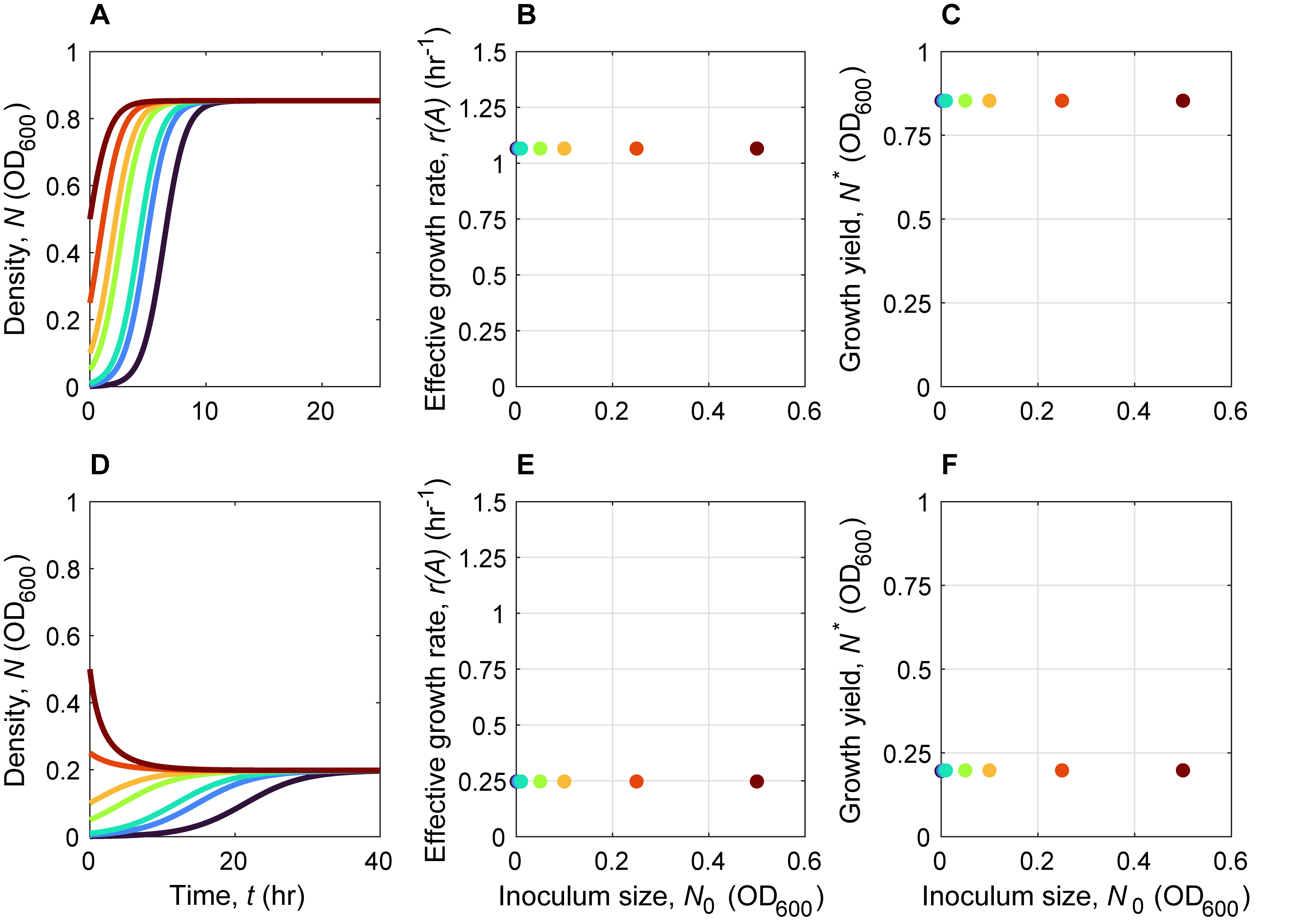

### S3 Fig

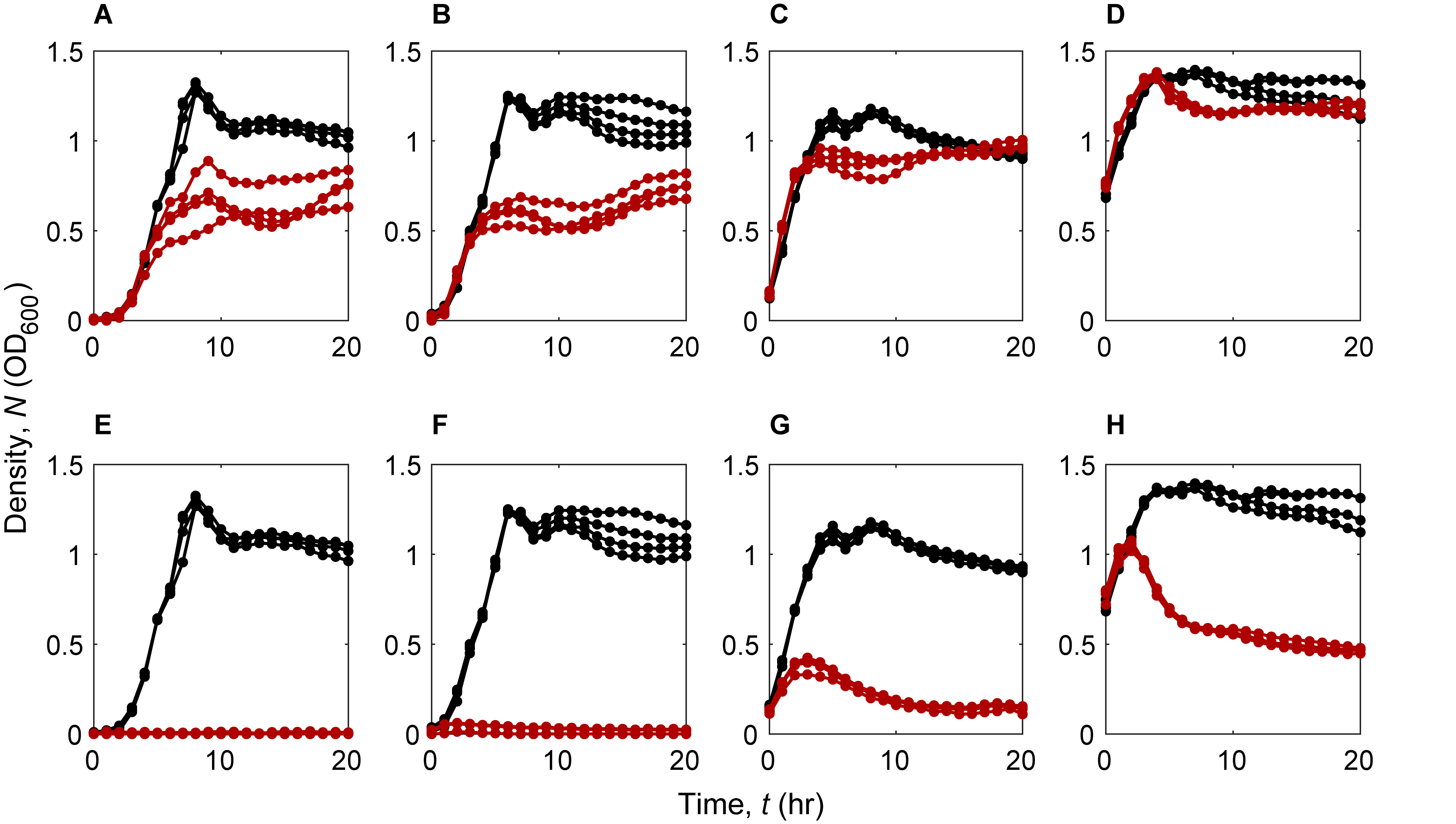

### S4 Fig

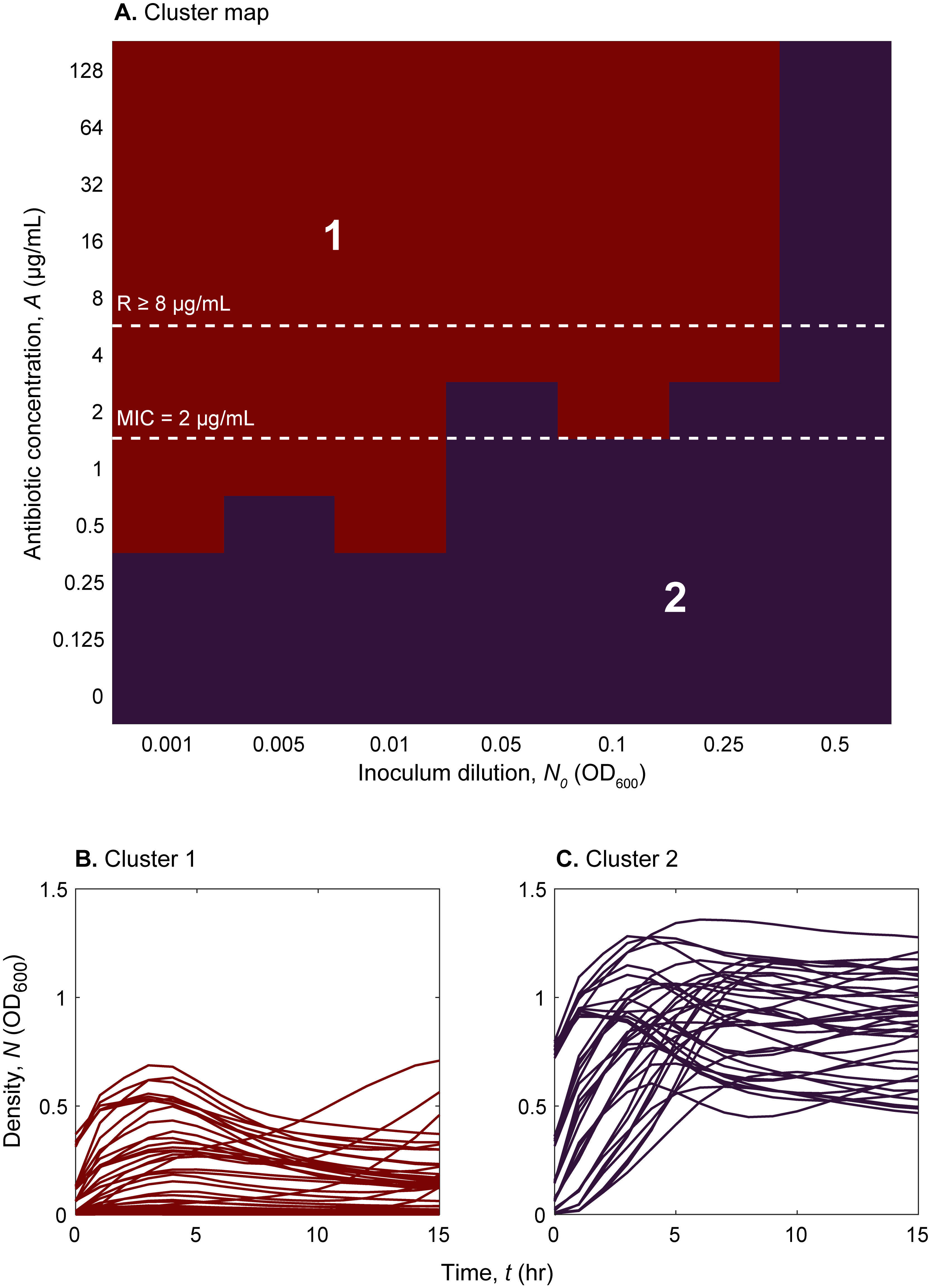

### S5 Fig

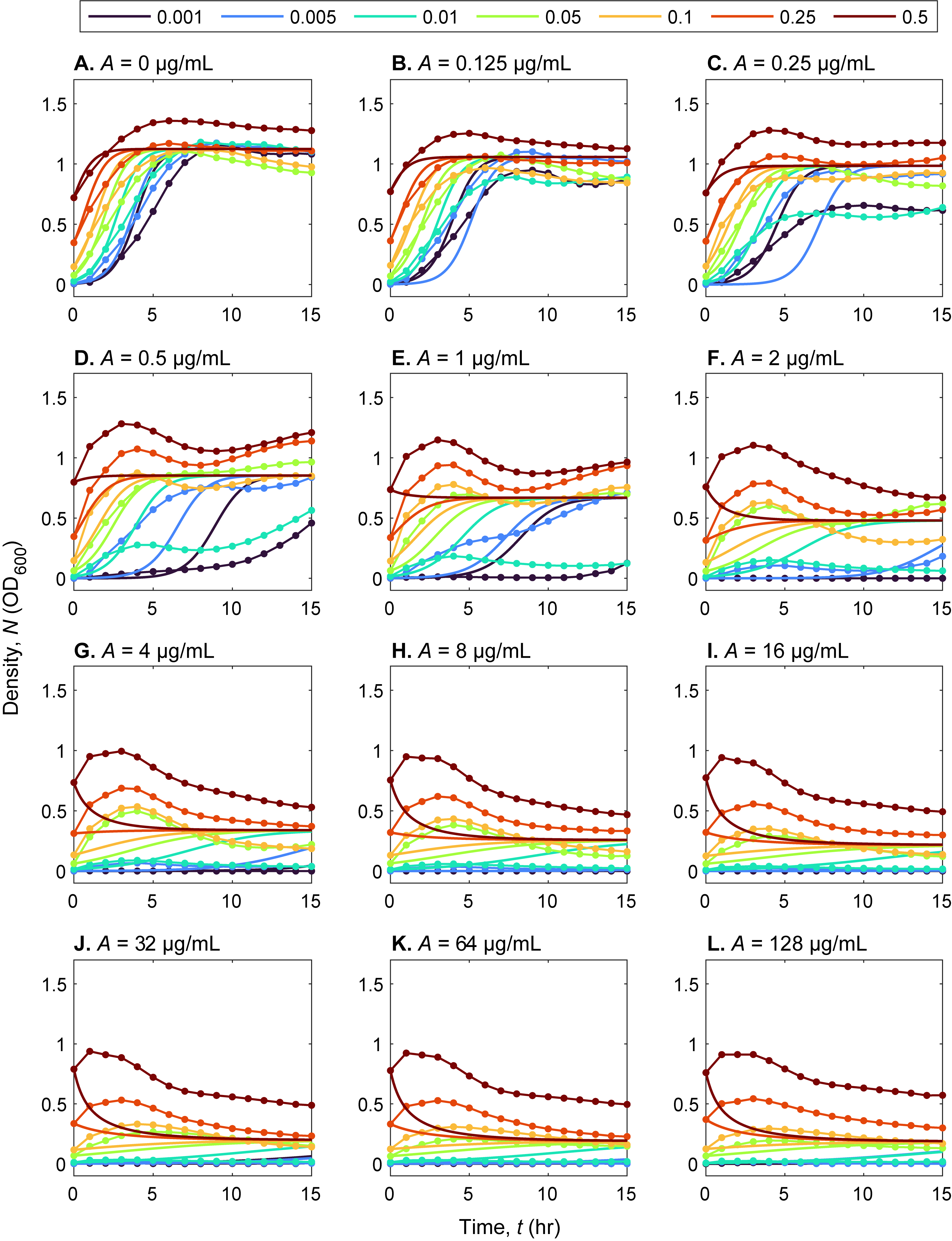

### S6 Fig

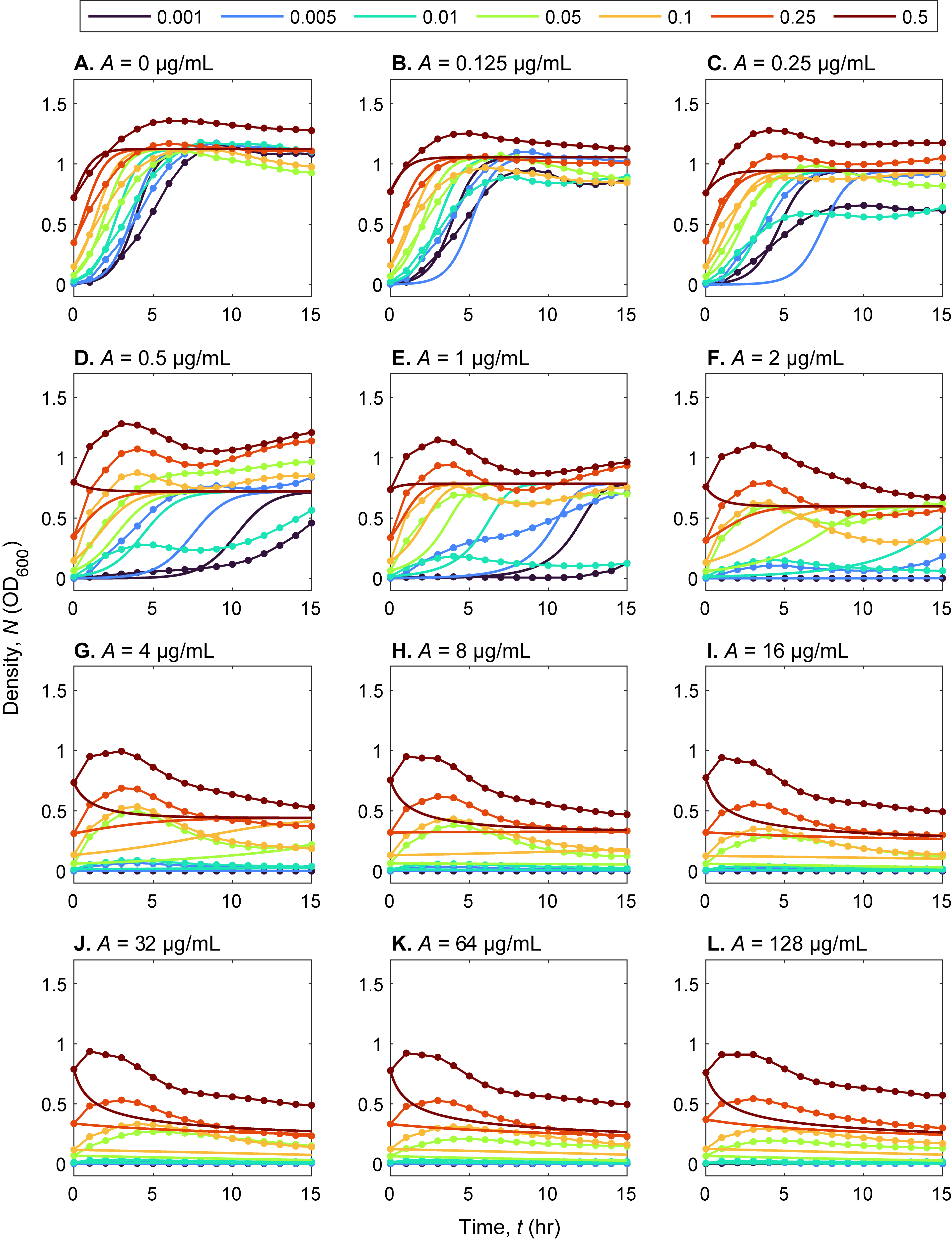

### S7 Fig

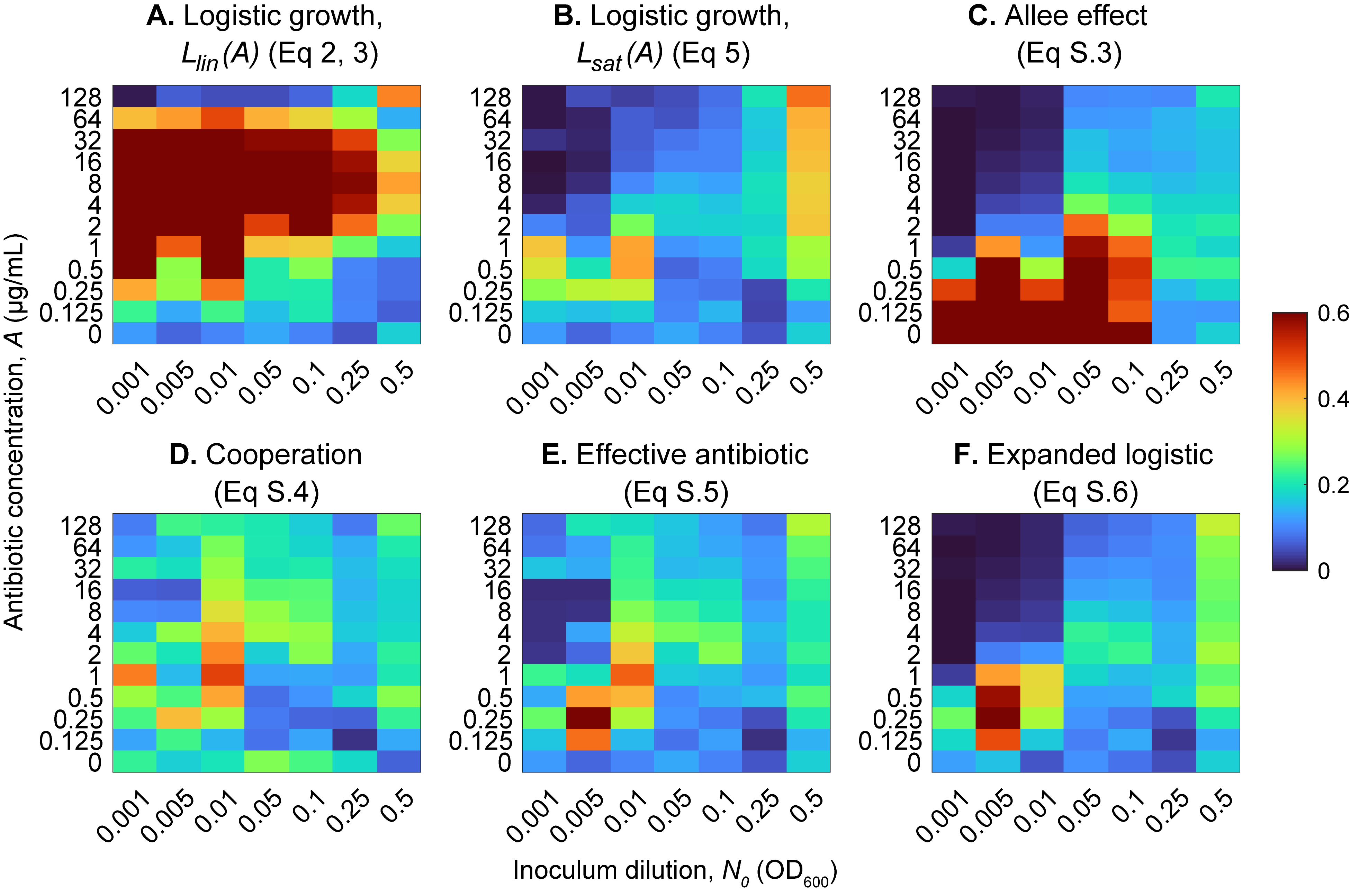
